# Asymmetry and ion selectivity properties of bacterial channel NaK mutants mimicking ionotropic glutamate receptors

**DOI:** 10.1101/2022.08.08.502946

**Authors:** Sonja Minniberger, Saeid Abdolvand, Sebastian Braunbeck, Han Sun, Andrew J.R. Plested

**Author notes:** Equally contributing authors (Sonja Minniberger, Saeid Abdolvand).

## Abstract

Ionotropic glutamate receptors are ligand-gated cation channels that play essential roles in the excitatory synaptic transmission throughout the central nervous system. A number of open-pore structures of α-amino-3-hydroxy-5-methyl-4-isoxazolepropionic-acid (AMPA)-type glutamate receptors became recently available by cryo-electron microscopy (cryo-EM). These structures provide valuable insights into the conformation of the selectivity filter (SF), the part of the ion channel that determines the ion selectivity. Nonetheless, due to the moderate resolution of the cryo-EM structures, detailed information such as ion occupancy of monovalent and divalent cations as well as exact displacement of the side-chains in the SF is still missing. Here, in order to resolve high-resolution crystal structures of the AMPA SF in its open-state, we incorporated the partial SF sequence of the AMPA receptor into the bacterial tetrameric cation channel NaK, which served as a structural scaffold. We determined a series of X-ray structures of NaK-CDI, NaK-SDI and NaK-SELM mutants at 1.42-2.10 Å resolution, showing distinct ion occupation of different monovalent cations. Molecular dynamics (MD) simulations of NaK-CDI indicated the channel to be conductive to monovalent cations, which agrees well with our electrophysiology recordings. Moreover, unique structural asymmetry of the SF was revealed by the X-ray structures and MD simulations, implying its importance in ion non-selectivity of tetrameric cation channels.

## 1. Introduction

Ionotropic glutamate receptors (iGluRs) belong to a large group of tetrameric ion channels with a common pore architecture^1^. At synapses in the vertebrate central nervous system, glutamate binding to AMPA subtype iGluRs leads to opening of an integral cation-permeable ion channel^2^. Recent advances in structural biology have confirmed the molecular details of the ion conduction pathway but the mechanisms behind ion selectivity, and in particular, non-selectivity between cations, remain obscure. Several open-pore structures of the AMPA-type glutamate receptor GluA2 in complex with auxiliary proteins are now available^3-5^. However, the resolution of cryo-EM maps is generally too low to unequivocally place ions in the pathway. Side chain orientations and ion binding sites thus remain vague. Our recent computational study found a monovalent cation single binding site^6^ at the same height as a presumptive hydrated Na^+^ ion adjacent to Q586 at the tip of the selectivity filter in the open-state homomeric GluA2 in complex with stargazin^4^.

Concepts of ion selectivity have been extensively developed on the prototypical K^+^-selective channel KcsA, in the light of structural studies^7,8^. Key features of this channel, the first to be crystallized, are two transmembrane spanning helices and a reentrant loop that forms the selectivity filter. This motif is shared among the large superfamily of tetrameric and pseudo-tetrameric ion channels, comprising highly selective, semi-selective and non-selective members. With the discovery of the non-selective cation channel NaK from *Bacillus cereus*, it was possible to identify key mechanistic differences between highly selective and non-selective channels^9-14^. Consistent with pre-structural predictions that ions should not pass independently through the pore^15^ in KcsA or NaK2K (a double mutant of NaK that is K^+^ selective), four contiguous K^+^ ion binding sites, formed by backbone carbonyl oxygens, coordinate along the full length of the selectivity filter. In contrast, there are only two distinct binding sites in NaK. Instead, a large water-filled vestibule replaces the other two binding sites and coordination of ions is only achieved by water molecules. The reduced number of binding sites and the presence of water inside the selectivity filter enhance flexibility and decrease the energetic barriers posed by desolvation of ions. Hence, the conduction mechanism and hydration state for Na^+^ or K^+^ are distinct from K^+^-selective channels.

Almost all ion channel pores are formed from oligomers, which may exist either in homomeric or heteromeric forms. Due to the heterogeneity of the sequence, heteromeric ion channels are obliged to be asymmetric. A number of experimental and computational studies revealed that asymmetric conformational changes in heteromeric ion channels may strongly influence ion permeation, selectivity, and gating^16.17^. Notably, most native AMPA receptors are heteromers^18,19^. Even in homomeric ion channels, structural asymmetry has been more frequently identified in recent studies using high-resolution X-ray analysis^20^, cryo-EM microscopy without applying symmetry averaging^21,22^ as well as NMR spectroscopy conducted under native condition^23,24^. Nevertheless, how exactly these asymmetric conformational changes in homomeric ion channel are correlated with the biological functions remains mostly elusive.

In our previous study of the wild-type NaK channel using a combination of solid-state NMR and molecular dynamics (MD) simulations, we observed a symmetry break in the selectivity filter (SF) in the Na^+^ conducting state^23^. Furthermore, the asymmetric conformational changes in the SF open up an unexpected side-entry ion pathway for Na^+^ permeation, which was not predicted before in other simulations starting from symmetric crystal structures. This ion permeation mechanism was supported by the recent high-resolution X-ray structure of NaK, revealing Na^+^ coordination at the side-entry ion binding site and structural asymmetry in the SF^20^. In another example from eukaryotic channels, the cryo-EM structure of the non-selective cation channel TRPV2 suggested a symmetry break from fourfold to twofold in the SF^22^.

In the present study, we substituted a part of the SF sequence from ionotropic glutamate receptors into the bacterial NaK channel. This approach was used before to investigate the ion permeation and selectivity of non-selective CNG channels^25,26^. We determined a number of high-resolution X-ray structures of the NaK-based glutamate receptor mimics (1.42-2.10 Å), showing distinct ion occupation of different monovalent cations in the SF. Using a MD based computational electrophysiology approach, we showed that the presented X-ray structure is mostly conductive to K^+^, and less conductive to Na^+^. Together with umbrella sampling simulations, symmetric SF conformation of the lower part of the SF was determined to be the main energetic barrier for efficient Na^+^ conduction. Electrophysiology recordings of the NaK-CDI mutant confirmed the channel to be conductive to monovalent cations. Finally, a unique structural asymmetry was determined in the NaK-based glutamate receptor mimics, which is most substantial in the mutated SF part. Together with the results from the MD simulations, we propose that structural asymmetry in the SF as a structural determinant of ion non-selectivity in tetrameric cation channels.

## 2. Results

### 2.1 Mimicking the glutamate channel pores using bacterial NaK as structural scaffold

The amino acid sequence of the SF in GluA2 does not resemble any other tetrameric ion channel and is relatively different to NaK (Fig. 1A). While there are some conserved residues in the pore helix M2, the selectivity filters only share a conserved glycine in the center of the filter. Also, the sequence is one residue longer in the NaK channel. We failed to incorporate the entire selectivity filter sequence of GluA2 (MQQCGI) into NaK. Accordingly, only parts of the filter could be mutated in the NaK channel to mimic the pore of GluA2, that is changing DGNF to C-DI. Mutations in the selectivity filter of NaKΔ19, did not alter the overall structure in NaKΔ18 C-DI. In order to mimic kainate receptors, DGNF was mutated to S-DI and also S-ELM. Altering positions equivalent to the Q/R site of GluA2, abolished assembly of the NaK channel, suggesting the crucial role of the TTV sequence in the NaK channel. To further assess the importance of the exchanged residues, the inverse mutations stemming from NaK were inserted into GluA2: MQQGCDI to TTVGCDI (GluA2-TTV) and MQQGDGNF (GluA2-DGNF) (Fig. 1A). The electrophysiological properties of these mutants are discussed below.

**Figure 1.**
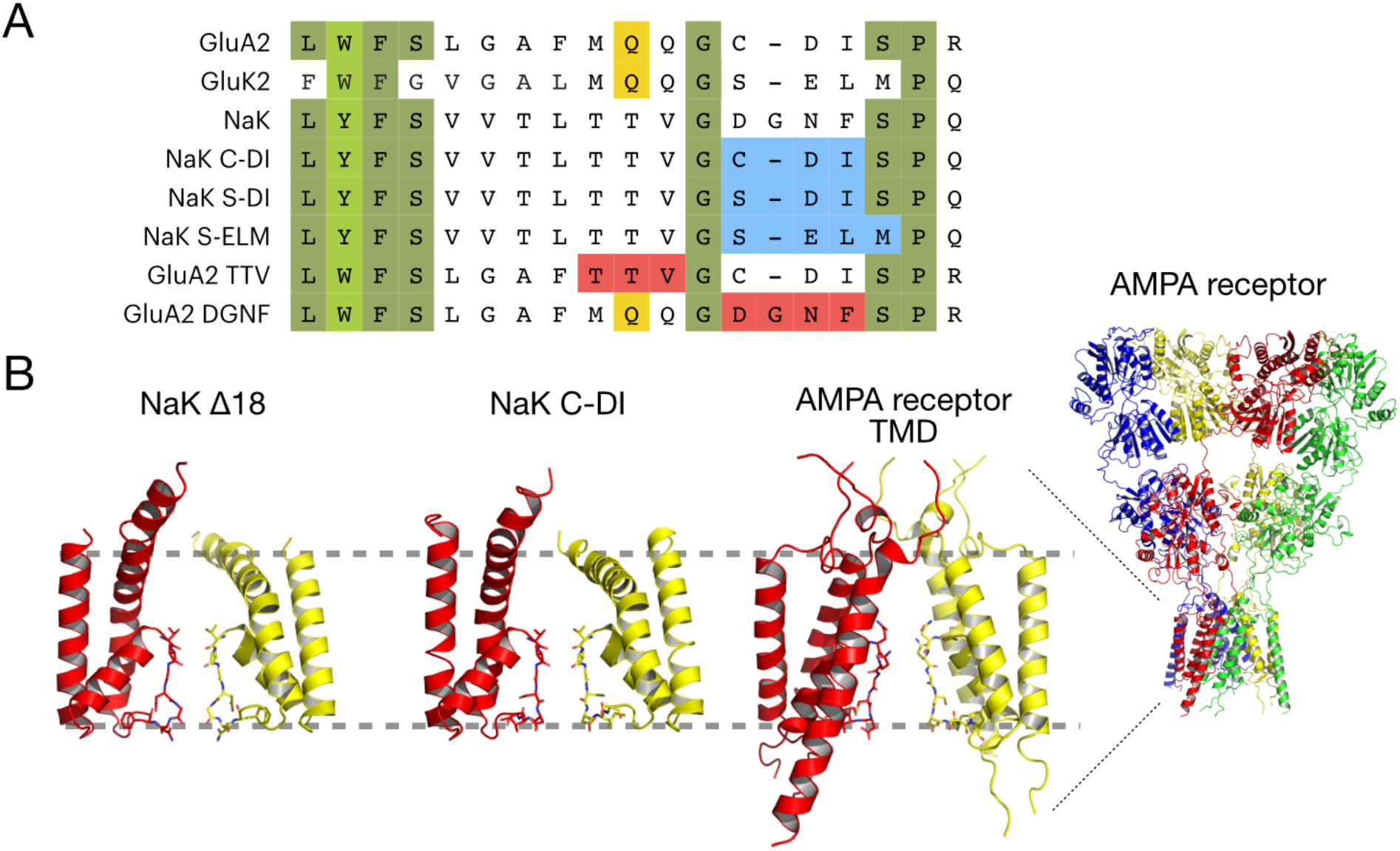
The non-selective channels NaK, NaK C-DI and GluA2. (A) Sequence alignment of the “pore loop” region of GluA2 with NaK and mutants of the NaK channel presented in this paper. Identical and similar residues are highlighted in dark and light green, respectively. The Q/R site is in yellow, while mutations introduced to NaK are highlighted in blue. Reverse mutations in GluA2 for patch-clamp recordings, adding mutations from NaK, are highlighted in red. (B) Comparison of the transmembrane domains of NaKΔ19 (PDB ID: 3E8H) and NaKΔ19 C-DI (PDB ID: 7OOR), both rotated 180°, with a zoom of the TMD of GluA2. Only two diagonally opposed subunits are shown for clarity. The extremes of the membrane are indicated with grey dashed lines. The entire AMPA receptor cryo-EM structure with auxiliary subunits removed (PDB ID: 5WEO) is shown on the right.

In comparison to NaK, GluA2 has an inverted structure with the selectivity filter on the intracellular side. When comparing to the glutamate receptor architecture, NaK lacks both the large extracellular domains as well as the third transmembrane helix (Fig. 1B).

### 2.2 Crystal structures of the NaK mutant C-DI

We determined 5 crystal structures of NaKΔ18 C-DI in the presence of different mixtures of monovalent ions (Na^+^, K^+^, Rb^+^, Li^+^ and Cs^+^, Fig. 2A-D) by molecular replacement using chain A of the NaKΔ19-K^+^ complex as a search model (PDB: 3E86) with the SF residues omitted. Following previous structural and functional studies on NaK, we crystallised both wild-type and the F92A mutant of the channel, because F92A showed an increase in ion conductivity compared to wild-type^10^. The structures were refined to resolutions between 1.47 Å to 2.10 Å. The highest resolution was obtained for the Na+ and K+ complex. All crystals except for the Cs+-bound structure were of space group C222_1_ while the latter crystallised in space group I4. The asymmetric unit contained two subunits from either one or two ion channel tetramers for C222_1_ and I4, respectively.

**Figure 2.**
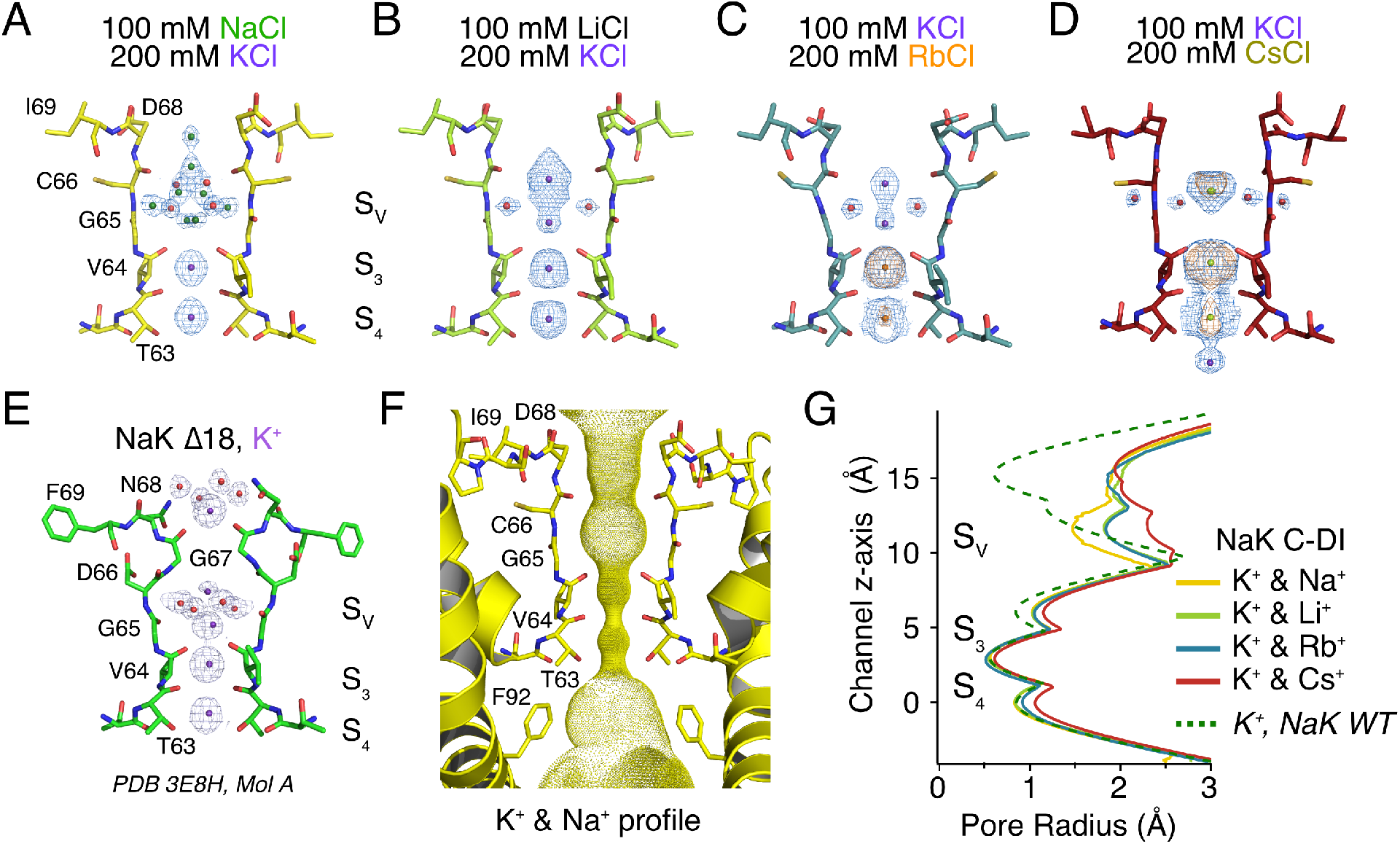
Monovalent ion binding in NaKΔ18 C-DI. (A-D) The 2Fo-Fc electron density maps (blue mesh), contoured at 1σ, show ion binding in the selectivity filter of NaKΔ18 C-DI. (A) Na^+^ (green) and water molecules (red) are present in the vestibule while K^+^ (purple) sits in binding sites S3 and S4. (B) Li^+^ contains too few electrons to contribute substantially to X-ray scattering at this resolution and cannot be seen. K^+^ and water molecules were therefore assigned to densities. Rb^+^ (C, orange) and Cs^+^ (D, olive) complexes were also solved, on a background of 100 mM KCl. Anomalous difference density for each ion species is shown in orange mesh, contoured at 3σ. (E) K^+^ complex of NaKΔ19 (PDB ID: 3E8H) for comparison. (F) Solvent-accessible pathway of NaKΔ18 C-DI generated with the program HOLE^27^. (G) Pore profiles of the different ion complexes of NaKΔ18 C-DI and NaKΔ19, generated with HOLE. The origin is set to the plane of hydroxyl oxygen of T63.

The SF of NaKΔ18 C-DI is one residue shorter than in NaKΔ19 and therefore straightens up, removing the upper restriction at the extracellular exit of the permeation pathway. The structure of the non-mutated motif TTV in the lower half of the SF was not altered by the mutations in the upper half. Calculation of the pore dimension using the program HOLE^27^ suggests that the major restriction site in NaK C-DI is the TTV part of the SF (Fig. 2F, G). Comparison of the SF structures of GluA2, wt-NaK and NaK C-DI revealed that GluA2 is wider than wt-NaK as well as NaK C-DI (Supplementary Fig. 1). Nevertheless, the side-chains of NaK CDI in the NaK C-DI mutant show similar orientation to the ones in the cryo-EM structures of GluA2 (Supplementary Fig. 1).

The differences in architecture between wt-NaK and the NaK C-DI mutant led to differences in ion distribution within the SF. In both channels, K^+^, Rb^+^ and Cs^+^ all occupy the same sites S_3_ and S_4_ (Fig. 2A, C and D). In contrast, no Na^+^ density was observed at these two sites in NaK C-DI, which is different to the previous result on wt-NaK^10^. In the vestibule, ion distribution differed between ion types present as well as compared to wt-NaK. In the alkali metal series, larger ions tended to have a stronger binding affinity in the SF (Fig. 2C,D). In the X-ray structure of the NaK C-DI mutant complexed with Li^+^ and K^+^, K^+^ ions occupy all sites in the SF (S_3_, S_4_ and S_v_), while Rb^+^ replaced K^+^ at the S_3_ and S_4_ sites and Cs^+^ filled all sites in the structures of these soaked complexes. Even though we observed two K^+^ binding sites in the vestibule in the K^+^ and Li^+^ and K^+^ and Rb^+^ complexes, the coordinating water molecules as seen in wt-NaK were not present. On the other hand, a mix of Na^+^ and water molecules occupied the vestibule in the Na^+^ and K^+^ complexes (Fig. 2A). In the Cs^+^ complex, weak anomalous density in the vestibule was observed only in one of the two independent tetramers of the asymmetric unit. We modeled this density with a Cs^+^ ion at 16% occupancy (Supplementary Fig. 2). We did not observe the ion binding site at the external entrance in wt-NaK^10^ in any of the X-ray structures of NaK C-DI mutants.

### 2.3 Crystal structures of NaK S-DI and S-ELM mutants

For a comparison between GluA2 and kainate receptors, we next determined the crystal structures of NaKΔ18 S-DI and S-ELM (NaK2Kai) in the presence of Na^+^ and K^+^ to resolutions between 1.42 Å and 1.85 Å. The substitution of cysteine to serine in NaKΔ18 S-DI did not alter the appearance or dimensions of the SF itself (Fig. 3A, B). In contrast, exchanging aspartate with glutamate at the external entrance created a more restricted entrance vestibule in NaKΔ18 S-ELM (Fig. 3C). Two opposing glutamate residues point towards the ion channel pathway and directly participate in ion coordination (Fig. 3A). The additional mutations to leucine and methionine did not influence the ion pathway. In the cavity on the intracellular side of the selectivity filter, the aromatic ring of F92 was oriented towards the ion conduction pathway. An alternative conformation in chain B limited the pore diameter to roughly 5 Å compared to about 10 Å in the F92A mutant (Supplementary Fig. 3)^28^. In both S-DI and S-ELM channels K^+^ is bound in binding sites S_3_ and S_4_. The distribution in the vestibule consisted of a mix of K^+^, Na^+^ and water (Fig. 3A). Despite prolonged soaking in a buffer containing Na^+^, the single ion, K^+^ could not be replaced in binding sites S_3_ and S_4_ in NaKΔ18 S-DI (Supplementary Fig. 4).

**Figure 3.**
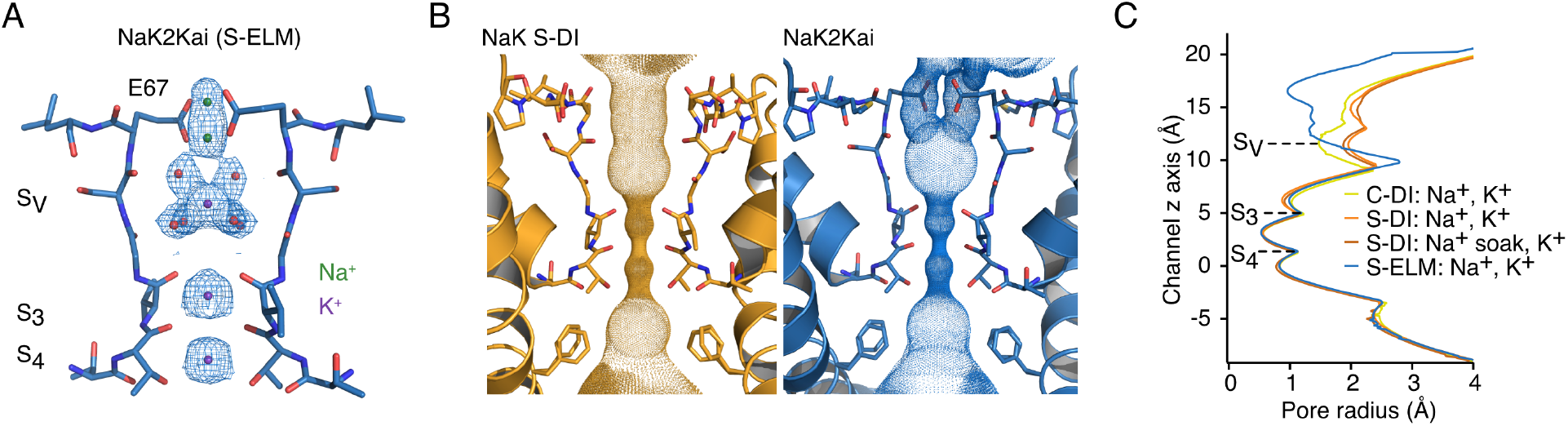
Structures of kainate receptor emulating mutants. (A) The NaK2Kai structure with Na^+^ and K^+^ in the SF. (B) Solvent-accessible surface of NaKΔ18 S-DI (left) and NaKΔ18 S-ELM (NaK2Kai, right) mutants. An alternative conformation of F92 can be observed in one of the two subunits of the asymmetric unit. (C) Pore profiles of the different NaKΔ18 mutants. Rotation of E67 towards the pore axis in the S-ELM mutant leads to a notable narrowing at the upper exit of the SF (above Sv), resembling the parent WT NaK channel.

### 2.4 Electrophysiology measurements

To investigate its ion permeation properties, NaK C-DI F92A was either purified and reconstituted into small unilamellar vesicles or in-vitro translated into nanodiscs. We recorded single-channel activity with the lipid bilayer technique in both mono- and bi-ionic conditions at different voltages. Only in one recording inward and outward permeation under bi-ionic conditions occurred while in the other recordings only mono-directional currents were observed independent of ionic composition (Fig. 4A). Single channel openings were in the range of 7.5 to 12 pA at -150 mV, giving rise to a single-channel conductance of 67 ± 10 pS. At +100 mV, single-channel conductance was 64 ± 6 pS, indicating no dependence of conductance on voltage. The overall conductance amounted to 66 ± 12 pS (Fig. 4B), which is substantially higher than the wild-type NaK conductance around 35 pS^29^. We further analysed single bursts of channel activity to determine open dwell-times. Three components were necessary to describe open dwell-times. The two fast components (35.3 ± 7.7 ms and 5.6 ± 0.3 ms) prevailed with a combined weight of 83% (Fig. 4C). Closed times required a fourth component in the exponential function to properly describe the life times. Overall, dwell times were longer in the closed state, with the fast components (62.6 ± 27.8 ms and 11.0 ± 6.3 ms) dominating. In contrast to the open state, there was a fourth, long component of over 2 s with a weight of 20 % (Fig. 4D). Dwell times are summarised in Supplementary table 2.

**Figure 4.**
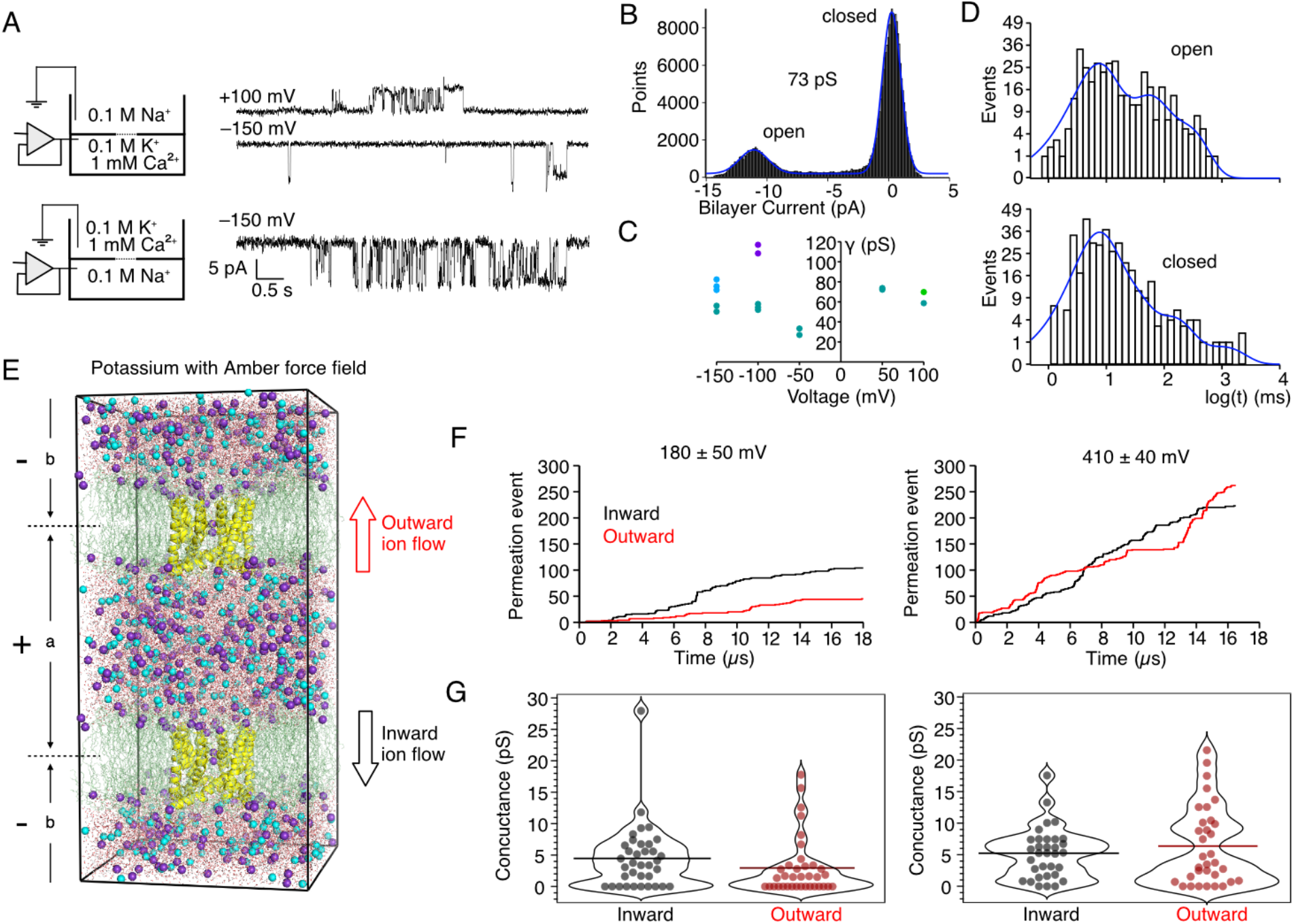
Conductance of NaKΔ18 C-DI. (A) Upper, Current traces of NaKΔ18 C-DI F92A fused from nanodiscs to a DPhPC bilayer. Representative traces at +100 mV and −150 mV. Lower, Current trace of NaKΔ18 C-DI F92A fused from *E. coli* total lipid extract proteoliposomes to a POPE:POPG bilayer. Openings were only seen at −150 mV. Buffer conditions in the two chambers of the bilayer apparatus are indicated in the schematics on the left. (B) All-points histogram of the unitary currents (observed in panel (A), lower trace) with the peak-fitting trace (blue) used to calculate conductance. (C) Conductance versus voltage, bionic conditions (Na^+^ vs K^+^ + Ca^2+^). Each colour represents one of four individual experiments, each circle is from one trace. (D) Exemplary open (upper) and closed (lower) dwell-time analysis of the lower current trace partially in panel (A). The histograms were fit with three and four exponential components, respectively (see Supplementary Table 2). (E) The double bilayer setup used for MD-based computational electrophysiology simulations including two NaKΔ18 C-DI proteins (yellow cartoon representation) embedded in two POPC lipid bilayers (green lines) surrounded by water molecules (red stick), K^+^ (purple balls) and Cl^−^ (cyan balls). Charge imbalances between the two compartments (a and b) corresponding to transmembrane voltages of 180 ± 50 mV and 410 ± 40 mV were maintained during the simulations. (F) Cumulative inward (black) and outward (red) ion permeations versus time for K^+^ simulations at a transmembrane voltage of 180 ± 50 mV and 410 ± 40 mV, respectively. The simulations were performed with AMBER99SB force field^30^. (G) Inward (black) and outward (red) K^+^ conductance for each 500 ns simulation run at transmembrane voltages of 180 ± 50 mV and 410 ± 40 mV. Mean of the conductance is shown as a horizontal line. The simulations were performed with AMBER99SB force field^30^.

The inverse mutations in GluA2 were tested with patch-clamp electrophysiology. We expressed GluA2 wt, GluA2-TTV and GluA2-DGNF in HEK-293 cells and exposed outside-out patches to 10 mM glutamate repeatedly. While GluA2-DGNF behaved similarly to GluA2 wt, GluA2-TTV remained unresponsive to glutamate even in the presence of desensitisation blocker CTZ (Supplementary Fig. 5), suggesting that it is either not folded, not conductive or not expressed at the membrane.

### 2.5 Molecular dynamics simulations of K^+^ permeation

Starting from the newly resolved crystal structures of NaK C-DI, we employed molecular dynamics based computational electrophysiology^31^ to simulate ion permeation in the channel. Using this method, an ion imbalance between the extracellular and intracellular sides of the channel remains during the simulations, leading to a constant transmembrane voltage that is similar to electrophysiology measurements (Fig. 4E). From the computational electrophysiology simulations, both inward and outward ion permeation through the channel can be observed simultaneously from one simulation setup.

We started the simulations with the crystal structure of NaK C-DI in the presence of Rb^+^ (PDB ID: 8A35), because this structure was the first crystal structure of the series that we solved. The simulations were conducted with 600 mM KCl at a transmembrane voltage of 180-430 mV. We performed simulations for a total time of 34.5 μs and 20 μs for the K^+^ simulations with AMBER99sb^30^ and CHARMM36m^32^ force fields, respectively. While comparable conductance of inward and outward K^+^ permeation was determined from the simulations with AMBER force field (Fig. 4F&G), we observed mostly inward K^+^ permeation during the CHARMM simulations (Supplementary Fig. 6). Averaged conductance of K^+^ in the AMBER simulations was ranging from 2.9 pS to 6.4 pS (Fig. 4F&G, Supplementary Table 3), which are substantially higher than the ones of wt-NaK (e.g.: 0.6 pS for inward K^+^, AMBER99sb force field) using the same method^23^.

During K^+^ permeation, sites S_3_ and S_4_ in the lower part of the SF formed by T63 and V64 are mostly occupied by K^+^ (Fig. 5). K^+^ occupancies at the same sites are also clearly visible in the crystal structures (Fig. 2A). Two additional weaker and more flexible K^+^ binding sites were observed in the vestibule of the SF that is filled with water molecules during the simulations. These results agree very well with the crystal structures of K^+^ & Li^+^ as well as Rb^+^ & K^+^ (Fig. 2B,C), revealing two K^+^ binding sites at nearly the same position. Furthermore, due to the exchange of one neutral residue N68 in wt-NaK by the negatively-charged Aspartic acid (D68) in the NaK C-DI mutant, several K+ were attracted at the upper mouth of the SF (Fig. 5). This observation is in contrast to the crystal structure, where no resolved electron density is visible at this region.

**Figure 5.**
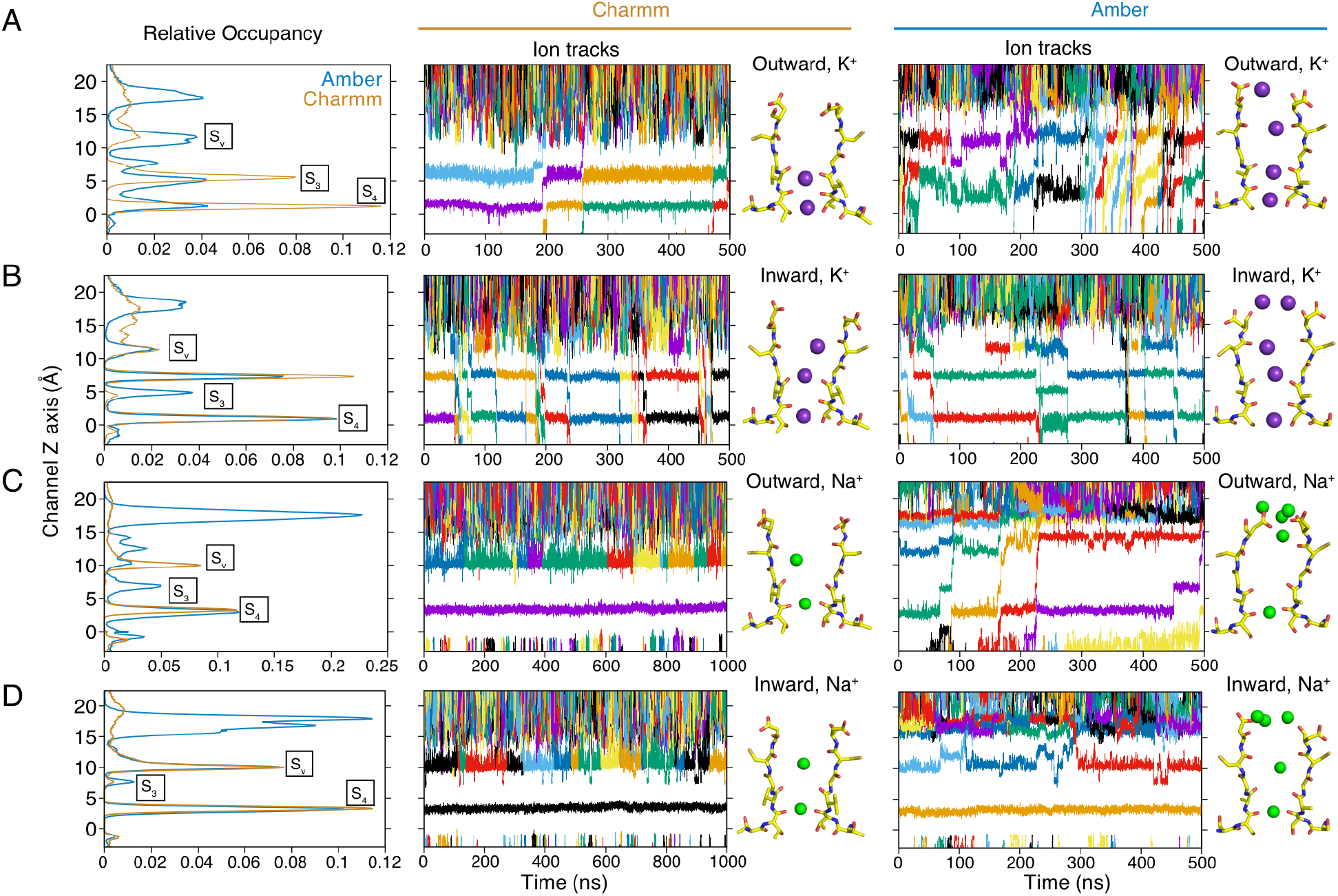
Molecular dynamics of ion permeation in NaKΔ18 C-DI. (A) For an outward K^+^ ion gradient, K^+^ occupancies in the selectivity filter (S3 and S4 at the lower part of the SF, Sv at the vestibule) are aligned with corresponding example ion track plots in the selectivity filter during the permeation and snapshots from the corresponding simulation trajectories for simulations performed with CHARMM36m^32^ and AMBER99SB^31^, respectively. K^+^ are shown as purple spheres. (B) as for (A) but with inward K^+^ gradient. (C) as for (A) but for Na^+^ permeation with an outward gradient. Note that in Charmm no Na^+^ permeation events occurred. Na^+^ are shown as green spheres in the snapshots. (D) As for (C) but for inward Na^+^ gradient - no permeation events occurred in either force field.

In both simulations at lower (180 ± 50 mV) and higher (410 ± 40 mV) transmembrane potentials, significant water co-permeations were observed. While the direction of the netto water flux corresponded to the direction of the ion permeation, the net number of water permeation was considerably lower than ion permeations (Fig. 6A). When analysing the water profile along the SF, a significant portion of water molecules were found in the vestibule, while water was nearly absent at S_3_ and S_4_ (Fig. 6B). Correspondingly, K^+^ were principally coordinated by backbone oxygens of T63 and V64, while the contribution of water molecules increased substantially in the vestibule and surpassed the protein oxygens (Fig. 6B). In the calculation of the number of ion-coordinating oxygens from both water molecules and protein residues, we considered only the first hydration shell, which is 3.4 Å for K^+ 6,33^.

**Figure 6.**
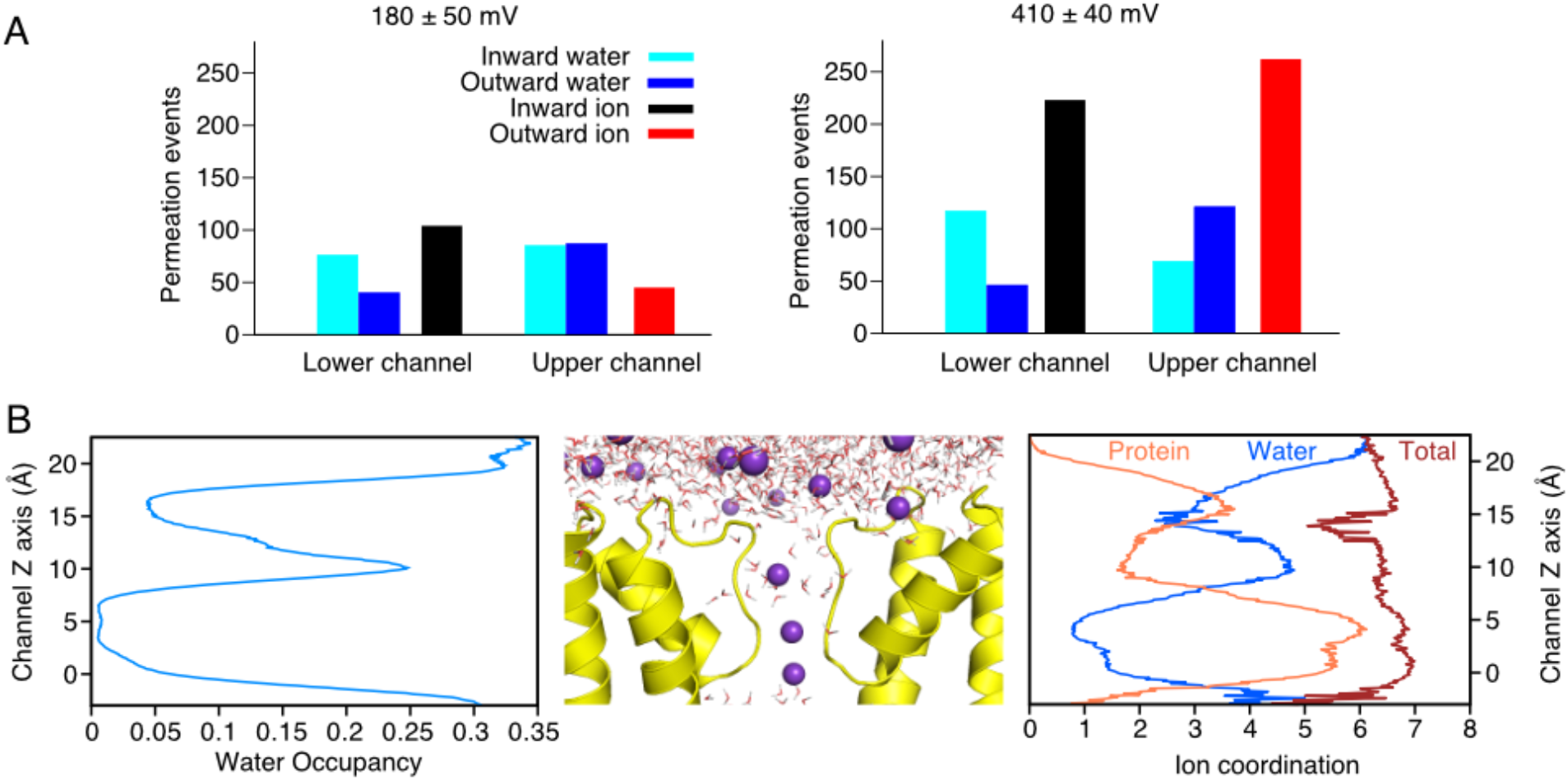
Water co-permeation. (A,B) Number of water molecules that permeated through the SF during inward (cyan) and outward (blue) K^+^ permeation. For comparison, the numbers of inward (black) and outward (red) K^+^ permeation are shown. The simulations were performed at a transmembrane voltage of 180 ± 50 mV and 410 ± 40 mV, respectively. (B, *left panel*) Percentage of water occupancy in and around the selectivity filter region. *(middle)* A snapshot selected from a representative simulation shows water molecules (depicted in stick) and K^+^ (purple spheres) in the SF. (*right*) The number of oxygens within the first hydration shell in the SF region along the pore axis. The number of coordinating water oxygens is in blue, the number of coordinating protein oxygens is in orange, and their sum is in red.

Additionally, we divided each K^+^ simulation run into 5 sub-trajectories and calculated the instantaneous conductance within each 100 ns. The instantaneous conductance was plotted against the distance between two F91 of opposing subunits in the M3 helix (d0, Supplementary Fig. 7), as well as the distances between two opposing residues in the SF (d1: distance between two side-chain oxygens of T63, d2: distance between two backbone oxygens of T63). Results shown in Supplementary Fig. 7 revealed no clear correlation between the conductance and the opening of the gates at the lower helix and the SF.

### 2.6 Molecular dynamics simulations of Na^+^ permeation

We also performed simulations of Na^+^ permeation at two different voltages (180-200 mV and 410-430 mV) and with two different force fields (AMBER99sb and CHARMM36m). In strong contrast to K^+^ permeation, only 6 Na^+^ permeation events could be detected within a total simulation time of 17.5 μs (Supplementary Table 3). During most of the simulation runs, Na^+^ bound tightly in the plane formed by four backbone carbonyls of T62 and thus blocked the entire channel (Fig. 5C and D). The same observation was reported previously in simulations of wt-NaK^20,23^. In the current study, we cannot make a direct comparison of Na^+^ occupancy at S_3_ and S_4_ between crystal structures and simulations, because in the crystal structure with mixed ion conditions no Na^+^ was visible in these two sites (Fig. 2A and Supplementary Fig. 4). Nevertheless, in the X-ray structure of the Na^+^ and K^+^ complex, Na^+^ densities are visible in the vestibule. Indeed, during the simulations at both negative and positive transmembrane voltages, we always observed substantial Na^+^ occupancy in the vestibule (Fig. 5 C and D).

To identify the energy barriers for Na^+^ permeation using the crystal structure of NaK C-DI as structural model, we calculated the potential of mean force (PMF)^34^ of Na^+^ in the NaK C-DI channel using the umbrella sampling method with CHARMM36m force field. For this purpose, we used the weighted histogram analysis method (WHAM)^35^. After equilibration of the system, one Na^+^ was pulled through the SF of the channel in the axial (z) direction with a force constant of 500 kJ mol^-1^ nm^-2^. Histograms of all umbrella sampling windows overlapped well except for a few regions. These two regions are located between S_3_ and S_4_ in the SF as well as S_3_ and S_v_ (Supplementary Fig. 8). Applying a force constant of 750 and 1000 kJ mol^-1^ nm^-2^ did not improve the sampling over these two regions. The PMF of umbrella sampling simulations shown in Supplementary Fig. 8 illustrates that Na^+^ binds tightly at S_3_ and S_4_ with a substantial energy barrier (about 30 kJ) between them.

### 2.7 Conformational plasticity and structural asymmetry in the selectivity filter

Compared to most of the reported crystal structures of NaK, a unique feature of NaK C-DI, NaK S-DI and NaK S-ELM is their structural asymmetry despite being homotetrameric cation channels (Fig. 7A,B). In this study we used the *relative difference between two neighbouring subunits* (Fig. 7D) to quantify the structural asymmetry. The SF distances between two neighbouring carbonyls revealed clear differences (Fig. 7A,B). An overlay of the SFs of chain A and chain B revealed no major conformational rearrangements (Fig. 7C). To exclude the possibility of a simple rigid body displacement of the whole subunit, we measured the distance between each residue in the whole channel. The largest relative difference was clearly observed in the SF part and especially for the mutated residues S-DI (Fig. 7E). Furthermore, we compared the relative difference of the SF for different mutants (C-DI, S-DI and S-ELM) under different ionic conditions, showing that the NaK S-DI mutant exhibits the largest structural asymmetry (Fig. 7D). In all structures, the difference was largest in the mutated upper part of the SF, while the TTV motif almost exhibited fourfold symmetry. The asymmetric deformation of the mutated part of the channel makes the SF more rectangular compared to the fourfold symmetric cases.

**Figure 7.**
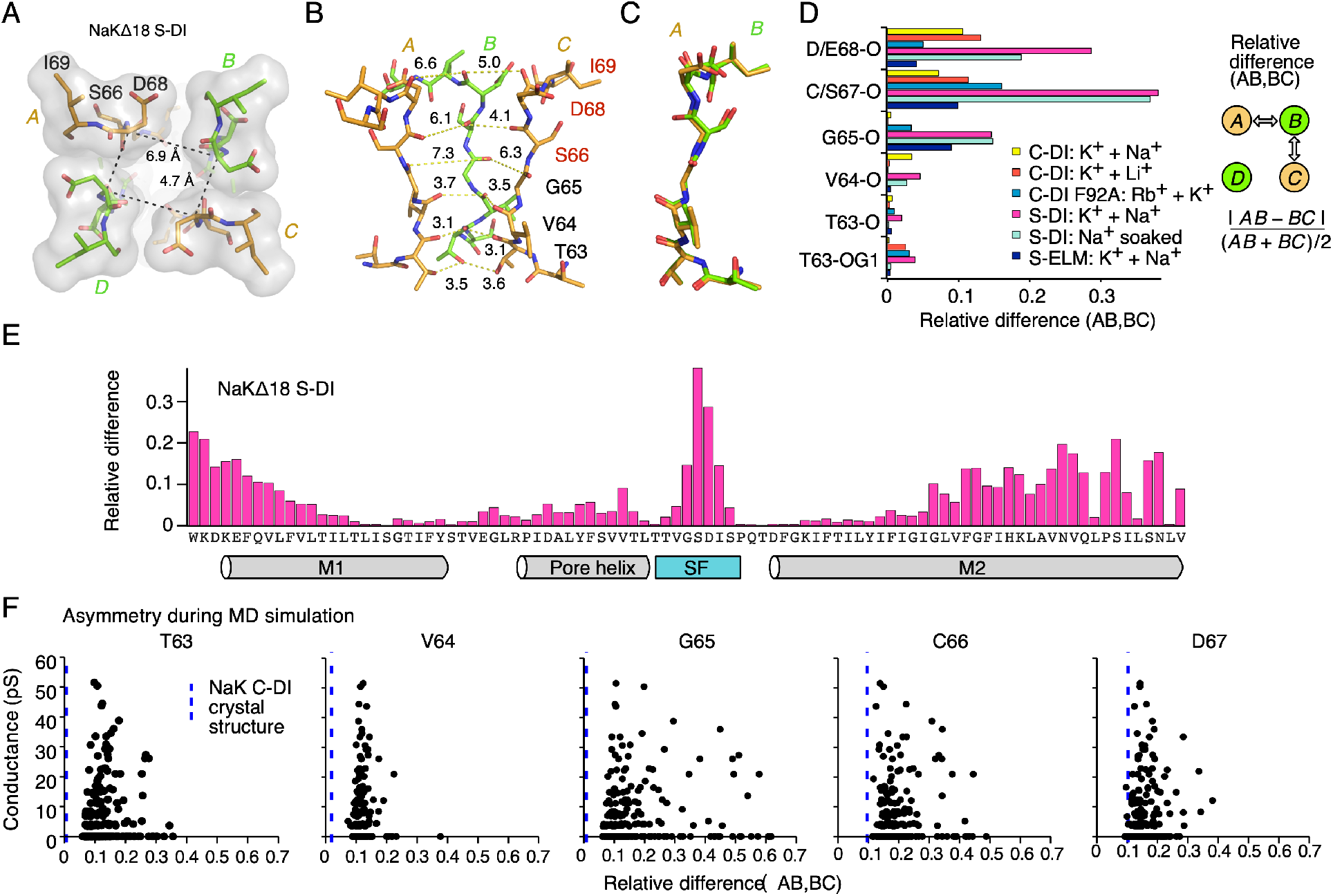
Structural asymmetry in X-ray structures of NaK mutants and during MD simulations. (A) Top view of NaKΔ18 S-DI. Symmetry-related subunits A and C, B and D, are coloured orange and green, respectively. The SF is shown in surface representation with the individual residues highlighted in stick representation. Distances between Cα-carbons of D68 between neighbouring subunits are indicated. (B) Side-view of the SF of NaKΔ18 S-DI with the same colouring as in (d). The front subunit was removed for clarity. Distances between oxygen carbonyls, as well as the side chain oxygen of T63, used for analysis, are indicated with dashed lines. (C) Structural alignment of subunit A with subunit B of NaKΔ18 S-DI. (D) Relative differences (rel. dif) between neighbouring side chain distances of NaKΔ18 mutants in different ionic conditions. A schematic representation of distances used for the relative difference parameter calculation. (E) Relative difference over the whole length of NaKΔ18 S-DI. The secondary structure elements are indicated underneath. (F) Simulated outward conductance as a function of the relative differece (AB,BC) parameter for the backbone oxygen atoms in the selectivity filter of the NaK C-DI. The simulations were performed with AMBER99SB force field.

In line with X-ray structures, MD simulations also revealed considerable flexibility and heterogeneity in the SF, especially in the mutated part as indicated by the root-mean-square-fluctuation analysis (RMSF, Supplementary Fig. 9). We calculated the relative differences between two subunits during the K^+^ and Na^+^ permeations for the NaK C-DI mutant (Supplementary Fig. 10). While Na^+^ and K^+^ simulations showed relatively similar distribution in their relative difference between two subunits of T63, the sampled conformations of all other residues are more asymmetric in the Na^+^ simulations compared to the K^+^ ones. Largest relative difference was observed for V64 in the Na^+^ simulations, while in the K^+^ simulations the mutated residues C66 and D67 in the upper part of the SF are more asymmetric. We further divided the K^+^ simulations into conductive and non-conductive states. Comparison of the d1-d4 distances between AB and CD subunits suggests the distances between two opposing subunits are significantly wider and more asymmetric in the K^+^ conductive simulations compared to the non-conductive runs (Supplementary Fig. 4E-H). The simulated conductance was finally plotted as a function of the relative difference (AB,BC) (Fig. 7F). Here it is interesting to note that the channel is most conductive with a certain range of asymmetry that is considerably larger than the one in the experimentally determined structures, while extreme large asymmetry again diminished the ion conductivity.

## 3. Discussions

In the present study we engineered a number of NaK channel mutants mimicking parts of the SF of ionotropic glutamate receptors. Unfortunately, we were only able to replace DGNF of the wt-NaK channel by C-DI, S-DI or S-ELM, but failed to change the TTV sequence of the SF to MQQ as in GluA2. In previous studies of the NaK based CNG mimics (NaK2CNG)^25,26^, due to the conserved sequence of the lower part of the SF, only DGNF sequence was altered. These results point to the crucial role of the TTV sequence in the correct assembly of the NaK channel. The same sequence could not be inserted into AMPA receptors to generate a functional channel. These observations indicate that although sequences and fold are conserved across the tetrameric channel superfamily, the dimensions of the selectivity filter region show variability and the accommodation of foreign elements requires geometric affordances that are not trivially implemented.

Our crystal structures are amongst the highest resolution structures of the NaK series to date. Recently, a comparable high-resolution structure (1.53 Å) of the wt-NaKΔ19 was published^20^. Interestingly, both high-resolution structures of NaK reveal structural asymmetry of the upper part of the SF, which was not observed before in any other X-ray structure of the NaK channel with lower resolution. These results raise the prospect that local asymmetry is more common in ion channels than previously anticipated and is revealed at high resolution. The functional roles of asymmetry are so far less clear, but from MD simulations we observed structure in the rates of permeation against geometry relating to degree of asymmetry.

In our ion binding analysis, we found distinct binding profiles of larger alkali ions (K^+^, Rb^+^ and Cs^+^) in the TTV part of the SF, while the larger vestibule at the upper part of the SF in NaK-CDI plays a much weaker role in binding and selectivity of ions. In line with this finding, our previous simulations^6^ and recent cryo-EM data^36^ of AMPA receptors revealed that the QQ part constitutes the major binding sites for monovalent ions, while the CDI part is wide and less important for coordination of monovalent ions with scarcely any metastable sites where ions reside. The present patch-clamp electrophysiology data showed that mutation of the CDI sequence in GluA2 by the DGNF of NaK had no obvious influence on channel activity, demonstrating again that the QQ filter in GluA2 is the most essential part of the SF for selection of monovalent ions.

Starting from the crystal structure of the NaK C-DI mutant, we performed a number of Na^+^ and K^+^ permeation simulations using AMBER99sb and CHARMM36m force fields. Both simulated and experimental K^+^ conductances of the NaK C-DI mutant are higher than the ones of wt-NaK, which might be attributed to the fact that the upper restriction at the extracellular exit of the permeation pathway is removed in the mutant structure compared to the wild-type one. Furthermore, consistent with our previous simulations of wt-NaK^23^, the X-ray structure is only conductive to K^+^, but essentially not to Na^+^ for which we observed only 6 permeation events across nearly 18 microseconds. Na^+^ bound tightly in the plane formed by four T63 or V64 carbonyls. Umbrella sampling of the Na^+^ permeation performed in the current study confirmed the high energy barrier between these two Na^+^ binding sites. Previous solid-state^23^ and solution-state NMR structures^24^ revealed that the lower part of the SF in the Na^+^-bound state is heterogeneous and less stable. In the present study, we were not able to determine the X-ray structures of NaK mutants containing Na^+^ solely. In all structures with Na^+^, K^+^ remain bound at sites S3 and S4. Therefore, we propose that the lower part of the SF with four-fold symmetry represents only the K^+^ stabilized state of the NaK mutant, which is not conductive to Na^+^. The lack of conductivity is almost certainly related in part to the deficiencies of the force field for Na^+^.

Ion occupancy, water coordination of ions within the SF, and the conformational states of the SF are highly similar for both simulation setups. Notably, we observe no direct correlation between K^+^ conductance and the opening of the gates at the lower helix and the SF. This is clearly different from previous simulation studies of potassium selective channels such as MthK and NaK2K, suggesting ion conductance can be enhanced by opening of the lower gate or the SF gate^28,37^.

From both crystal structures and MD simulations, we observed considerable structural asymmetry especially at the upper part of the SF. In the crystal structures of the NaK mutants, the upper part of the SF showed clear differences in the distances between two neighbouring subunits for C(S)66 and D68. Although the backbone and the side-chain conformation remain nearly unchanged for each chain, the upper part of the SF becomes substantially more rectangular compared to the fourfold symmetric NaK structures. In clear contrast to our finding, a recent crystal structure of wt-NaK also revealed conformational asymmetry for the corresponding residues D66 and N68, but with significant conformational changes in their backbone and side-chain^20^. Structural asymmetry of the SF in wt-NaK and NaK mutants observed in the X-ray structures are in line with recent solid-state^23^ and solution-state NMR^24^ studies on the wt-NaK, showing conformational heterogeneity and plasticity in the filter. Another example could be found in the eukaryotic system, where cryo-EM structure of the non-selective TRPV2 channel suggested a symmetry break from fourfold to twofold in the pore domain^22^. Considering these recent evidences of conformational asymmetry in different non-selective cation channels, our current study here again demonstrated the conformational asymmetry in the pore region as a unique feature for ion non-selectivity in tetrameric channels. We also noted that a number of recent X-ray structures of TREK-1 K2P channels showing asymmetric K^+^-dependent conformational transition in the SF that was suggested to be responsible for C-type gating^38^. All these examples demonstrated that ion selectivity and conductance can be regulated by local asymmetrical conformational changes, which is probably the energetically most efficient manner to alter ion channel function without requiring conformational transition of all subunits at large-scales. Native glutamate receptor ion channels are perhaps exclusively heterotetramers, giving a further level of local asymmetry. Our work shows that heteromerisation is not required for asymmetry and associated tuning of ion permeation.

## 4. Material and Methods

### 4.1 Online data availability

Data deposition: The atomic coordinates have been deposited in the Protein Data Bank, www.pdb.org [PDB ID codes 7OOR (C-DI Na^+^/K^+^), 7OOU (C-DI Li^+^/K^+^), 7PA0 (C-DI F92A K^+^/Rb^+^), 8A7X (C-DI F92A K^+^/Cs^+^), 7OPH (S-DI Na^+^/K^+^), 7OQ2 (S-DI Na^+^ soak), 7OQ1 (S-ELM Na^+^/K^+^), and 8A35 (C-DI Rb^+^/Na^+^)].

### 4.2 Methods

#### 4.2.1 Cloning and mutagenesis

The full-length NaK channel from *Bacillus cereus*^9^ was synthesised and cloned into the pQE60 expression vector using *NcoI* and *BamHI* restriction sites. A thrombin cleavage site was introduced between the C-terminus of the protein and a hexahistidine purification tag. To obtain open-channel structures the first 18 amino acids at the N-terminus were removed (NaKΔ18). This construct is termed Δ19^10^. All NaKΔ18 mutants are based on this construct.

#### 4.2.2 Expression and purification

Protein expression and purification were performed as described^10^. Briefly, expression in NEBExpress *I*^*q*^ *Escherichia coli* cells was induced with 0.4 mM IPTG (isopropyl-β-D-thiogalactoside) at 25 °C for 20 h. 5-10 mM BaCl_2_ were added to the expression medium for the NaKΔ18 C-DI F92A mutant. Cells were lysed in 50 mM Tris-HCl, pH 8.0, 100 mM of the respective salt (NaCl, KCl, LiCl, RbCl) and 1 mM phenylmethylsulfonyl fluoride and extracted with the addition of 40 mM n-decyl-β-D-maltoside (DM) for 3 h at room temperature. Extracted protein was purified on a Talon Co^2+^-affinity column. After elution the hexahistidine tag was removed either by incubation with 1 unit of thrombin per 2 mg protein overnight at room temperature or with 1:250 (molar ratio) trypsin for 1 h at room temperature. Finally, the protein was run on a Superdex 200 increase (10/300) column in 20 mM Tris-HCl (NaKΔ18 C-DI F92A, NaKΔ18 S-DI, NaKΔ18 S-ELM) pH 8.0 or HEPES (NaOH) (NaKΔ18 C-DI) pH 7.5, 5 mM DM and 100 mM of the corresponding salt (NaCl, KCl, LiCl, RbCl, CsCl).

#### 4.2.3 Protein crystallisation

Purified protein was concentrated to ∼12 mg/ml using an Amicon ultracentrifuge filtration device (50 kDa molecular weight cutoff) and crystallised using both sitting drop and hanging drop vapour diffusion method by mixing equal volumes of protein and reservoir solution. Well solution for NaKΔ18 C-DI crystals with NaCl or LiCl contained 40 % (v/v) (±)-2-methyl-2,4-pentanediol (MPD) and 200 mM KF or 200 mM K acetate, respectively. NaKΔ18 C-DI crystals with RbCl were grown over a well solution of 30 % MPD, 100 mM HEPES (NaOH) pH 7.5 and 5 % polyethylene glycol (PEG) 4000. NaKΔ18 S-DI crystals appeared over well solutions of 62 % MPD, 100 mM HEPES (KOH) pH 7.5 and 200 mM KCl, while NaKΔ18 S-ELM grew over 47 % MPD, 100 mM HEPES (NaOH) pH 7.5. Several crystals of NaKΔ18 S-DI were soaked in 100 mM NaCl, 100 mM Tris-HCl pH 8.0 and 50% (v/v) MPD. Finally, NaKΔ18 C-DI F92A crystals with CsCl were obtained with 63 % MPD, 100 mM Tris-HCl pH 8.0 and 200 mM CsCl and additionally soaked in 90 mM CsCl, 100 mM Kryptofix-222 and 50 % MPD. Rb^+^ complex crystals were grown over a well solution of 63 % (v/v) MPD and 100 mM Tris-HCl pH 8.0. Crystals were directly frozen in liquid nitrogen with the crystallisation solution serving as the cryoprotectant. Most crystals belonged to the space group C222_1_ with similar unit cell dimensions of around *a* = 81 Å, *b* = 88 Å, *c* = 49 Å, α = β = γ = 90°. The Cs^+^ crystal and the Rb^+^ - NaKΔ18 C-DI crystal belonged to space group I4 with unit cell dimensions of *a* = *b* = 68 Å, *c* = 90 Å, α = β = γ = 90°. The latter exhibited strong merohedral twinning with a twin fraction of around 44 %. The pseudo-fourfold axis of the channel tetramer coincided with the crystallographic fourfold- or twofold axis for space group I4 or C222_1_, respectively.

#### 4.2.4 Structure determination

NaKΔ18 C-DI and NaKΔ18 C-DI F92A X-ray diffraction data were collected at 100 K at beamline BL14.1 at BESSY II (Berlin, Germany)^39^. Anomalous data were collected for the Rb^+^ complexes. NaKΔ18 S-DI and NaKΔ18 S-ELM X-ray diffraction data were collected at beamline P11 at PETRA III at DESY (Hamburg, Germany)^40,41^. Data processing and scaling were performed using either XDSAPP^42^ or DIALS^43^ and the CCP4i suite^44^. All structures were solved by molecular replacement using chain A of NaKΔ19 (PDB 3E86) as a search model with the selectivity filter residues omitted and followed by repeated cycles of refinement in the Phenix suite^45^ and manual model building in Coot^46^. The quality of all structures was checked with PDB_REDO^47^. All structural figures were prepared using PyMOL () software. Pore profiles were calculated in the program HOLE^27^.

#### 4.2.5 MD Simulations

For studying the Na^+^ and K^+^ permeation through the NaK channel, we performed MD simulations with a total time of 44.5 μs for K^+^ and 30 μs for Na^+^ using the Groningen Machine for Chemical Simulations (GROMACS)^48^ software package version 5.1.2. In all simulations, the high-resolution crystal structure of NaK C-DI (with the F92A mutation introduced in silico) was embedded in a patch of a 1-palmitoyl-2oleoyl-sn-glycero-3phosphocholine (POPC) membrane and solvated in water with K^+^ or Na^+^ and Cl^-^ ions, corresponding to a salt concentration of 600 mM. The F92A mutant instead of wild-type (WT) NaK channel was used in all simulations to increase the currents^10^. All titratable residues of the protein were protonated according to their standard protononation state at pH 7. During the simulations, the Na^+^ and K^+^ ions were driven by the transmembrane voltage induced by charge imbalances in dual membrane setup as implemented in the Computational Electrophysiology (CompEl) method^31^ in GROMACS. Simulations are performed in both AMBER99sb^30^ and CHARMM36m^32^ force fields using TIP3P water model^49^.

For simulations with the AMBER99sb force field, Berger’s parameters for lipids^50^ and Joung’s parameters for ions^51^ were used for the equilibrium and production simulations. Using virtual sites instead of hydrogen bonds in protein, allowed for a 4 fs time steps for integration^52^. To equilibrate the systems, a short 10 ns simulation was performed by position-restraining all heavy atoms of the NaK channel with a force constant of 1000 kJ mol^-1^ nm^-2^ to the starting structure. The system was further equilibrated for 20 ns by releasing the position restraints. In production simulations, two different charge imbalances of 2e^-^ and 4e^-^ were applied for both Na^+^ and K^+^ ions which induced transmembrane voltages of ∼370 mV and ∼ 620 mV respectively.

For simulations with the CHARMM36m force field, the simulation setups, GROMACS inputs and parameters were provided by CHARMM-GUI webserver^53^. The equilibrations were performed using the standard GROMACS inputs of the CHARMM-GUI. The charge imbalances of 2e^-^ and 4e^-^ were applied for both Na^+^ and K^+^ ions. A time step of 2 fs was used for all simulations.

For all simulations, short-range electrostatic interactions were calculated with a cutoff of 1.0 nm, whereas long-range electrostatic interactions were treated by the particle mesh Ewald method^54,55^. The cutoff for van-der-Waals interactions was set to 1.0 nm. Simulations were performed at a constant temperature 310 K with a velocity rescaling thermostat^56^. The pressure was kept constant at 1 bar by means of a semi-isotropic Brendensen barostat^57^. All bonds were constrained with the LINCS algorithm^58^.

#### 4.2.6 Umbrella Sampling

The potential of mean force (PMF) was calculated for Na^+^ using umbrella sampling method^34^. For this purpose, an equilibrated single bilayer simulation setup with Na^+^ with Charmm force field was the starting point. A single Na^+^ ion was pulled along the central axis (z axis) of the selectivity filter of the NaK C-DI channel with a force constant of 500 kJ mol^-1^ nm^-2^. Based on the pulling simulation trajectory 73 umbrella windows were generated which provided a window spacing of 0.3 Å. For calculating the PMF, weighted histogram analysis method (WHAM)^35^ was used which is implemented in GROMACS^59^. In each umbrella sampling window, three different harmonic biasing potential with spring constants of 500, 750 and 1000 kJ mol^-1^ nm^-2^ were applied to hold the ion. The final PMF plot was computed based on the spring constant of 1000 kJ mol^-1^ nm^-2^. Each window was simulated for 0.5 ns.

#### 4.2.7 Electrophysiology-Patch-clamp

Rat GluA2 mutants were made by overlap PCR and expressed in HEK 293 cells by transient transfection^60^. The external solution (Ringer solution) contained 150 mM NaCl, 0.1 mM MgCl_2_, 0.1 mM CaCl_2_, 5mM HEPES pH 7.3, titrated with NaOH. A glutamate stock solution consisted of 2 M L-glutamic acid, 0.5 M D(+)saccharose was diluted with external solution to a final concentration of 10 mM glutamate for experiments. Glutamate was applied to outside out patches from HEK293 cells with a fast-perfusion tool^61^. For the GluA2 TTV mutant, the desensitisation blocker cyclothiazide (CTZ, 100 µM)^62^, was included in some experiments. In some experiments, a different buffer with, 158 mM NaCl, 3 mM KCl, 1 mM CaCl_2_, 20 mM HEPES pH 7.3, titrated with NaOH, was used. The pipette solution contained 115 mM NaCl, 1 mM MgCl_2_, 0.5 mM CaCl_2_, 10 mM NaF, 5 mM Na_4_BAPTA, 10 mM Na_2_ATP, 5 mM HEPES, titrated to a pH of 7.3 with NaOH. Patches were clamped at -80 mV to +80 mV. Currents were recorded using Axograph X (Axograph Scientific) via an Instrutech ITC-18 interface (HEKA) at 40kHz sampling rate and filtered at 10 kHz (−3 dB cutoff, 8-pole Bessel).

#### 4.2.8 Bilayer electrophysiology

NaKΔ18 C-DI F92A was used for single-channel recordings. Additionally, this construct was cloned into pEXP5-CT/TOPO expression vector for *in vitro* translation directly into 1,2-diphytanoyl-sn-glycero-3-phosphocholine (DPhPC) nanodiscs. The purified protein was reconstituted into either *E. coli* total lipid extract or POPE:POPG (7:3) (1-palmitoyl-2-oleoyl-sn-glycero-3-phosphoethanolamine: 1-palmitoyl-2-oleoyl-sn-glycero-3-phospho-(1’-rac-glycerol) (sodium salt)) proteoliposomes at a protein:lipid ratio of 0.1-20 µg per mg lipid. After dialysis against 500 mM KCl, 10 mM HEPES (KOH) pH 7.5, aliquots were stored at -80°C until use. Bilayer recordings were performed in a home-made chamber (gift from Chris Miller, Brandeis University, Waltham, MA) using an Elements SRL (Cesena, Italy) eONE amplifier. The bilayer was formed in the aperture of 50-80 µm diameter by the method of painting using DPhPC or POPE:POPG (7:3). Different buffer combinations for cis- and trans-chambers were used: 100 mM KCl, 10 mM HEPES (KOH) pH7.5 and 1 mM CaCl_2_, against either 100 mM NaCl or 100 mM KCl and 10 mM HEPES (NaOH or KOH, respectively) pH 7.5. In one case the cis-chamber was filled with 100 mM NaCl, 10 mM HEPES (NaOH) pH 7.5, 1 mM CaCl_2_, and the trans-chamber with 125 mM NaCl, 0.5 mM CaCl_2_, 10 mM NaF, 5 mM HEPES (NaOH) pH 7.2 and 0.05 mM spermine. Voltage across the bilayer was applied using Ag/AgCl electrodes immersed directly into each chamber. Bilayer formation was monitored applying a triangular voltage protocol from -100 mV to +100 mV, observing the rise in capacitance. After successful bilayer formation, NaKΔ18 C-DI F92A nanodiscs were added to the grounded cis-compartment whereas proteoliposomes were added to the trans-side. Fusion of channels with the bilayer were detected through observation of single channel currents. Currents were recorded at different holding potentials between -150 mV and +100 mV. Signal acquisition was done with the Elements Data Reader. Unfiltered traces were exported in Axon binary format, opened in Axograph X and low-pass filtered at 100 kHz. Bursts of single channel activity were further exported and dwell-time analysis was performed with ASCAM (Advanced Single Channel Analysis for Mac, available at github.com/AGPlested/ASCAM). Histograms of events were binned according to log-transformed values, and fitted with double-, triple- or quadruple-exponential equations, following the general functional form:

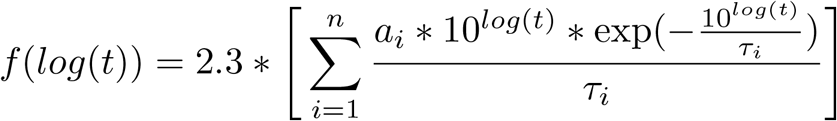

where *n* was 2, 3 or 4 respectively, *a*_i_ are the amplitudes of each component, and *τ*_i_ are the time constants.

## 5. Acknowledgment

This work was funded by the Deutsche Forschungsgemeinschaft (DFG) RU2518 DynIon to A.J.R.P. (P3; PL 619/5-1) and H.S. (P3; SU 997/1-1), the DFG under Germany’s Excellence Strategy—EXC 2008–390540038—UniSysCat to H.S., a DFG Heisenberg Professorship (PL 619/3-1 and PL619/7-1) to A.J.R.P., and the European Research Council Consolidator Grant “GluActive” (647895) to A.J.R.P. Diffraction data have been collected on BL14.1 at the BESSY II electron storage ring operated by the Helmholtz-Zentrum Berlin (Mueller et al., 2015). We would particularly like to acknowledge the help and support of Manfred Weiss and his team during the experiment. We acknowledge DESY (Hamburg, Germany), a member of the Helmholtz Association HGF, for the provision of experimental facilities. Parts of this research were carried out at PETRA III and we would like to thank Dr. Johanna Hakanpää for assistance in using beamline P11. The computations were performed with resources provided by the North-German Supercomputing Alliance (HLRN). We gratefully acknowledge the Gauss Centre for Supercomputing e.V. (https://www.gauss-centre.eu/) for funding this project by providing computing time through the John von Neumann Institute for Computing on the GCS Supercomputer JUWELS at Jülich Supercomputing Centre.

