## Supplementary Figures and Tables for "Asymmetry and ion selectivity properties of bacterial channel NaK mutants mimicking ionotropic glutamate receptors"

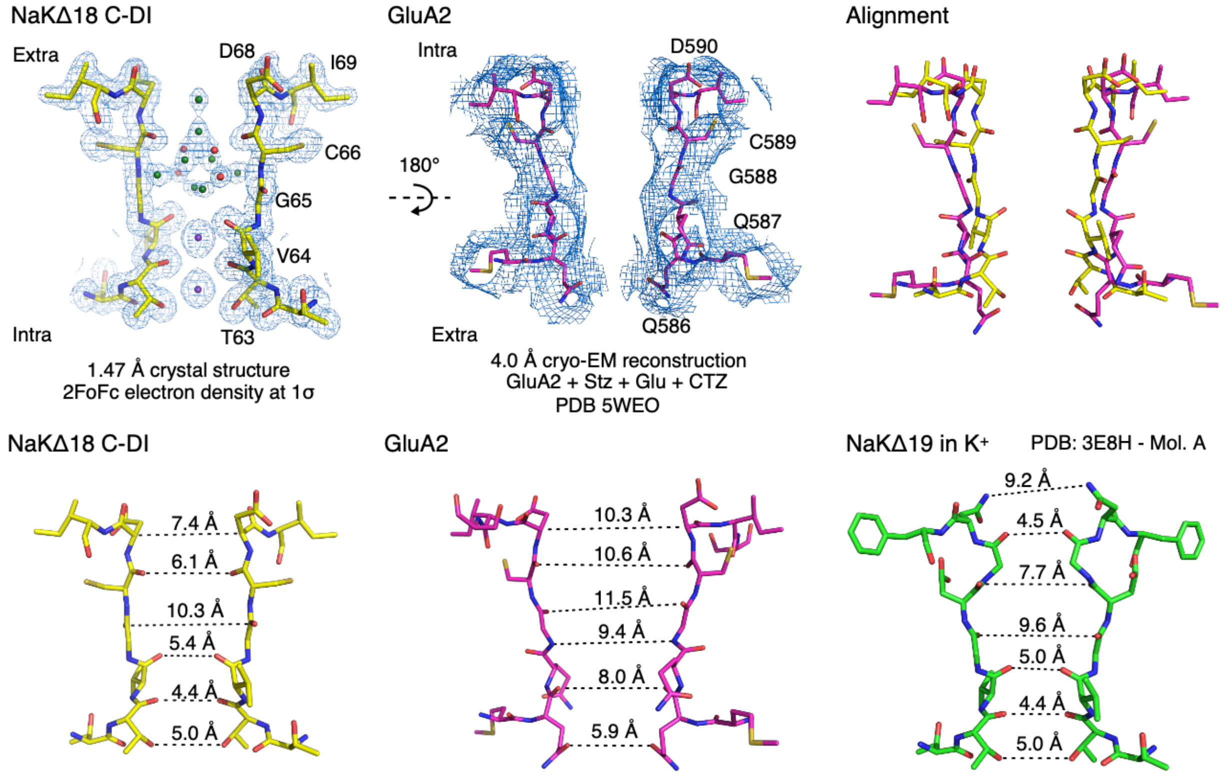


**Supplementary Figure 1 Comparison of NaKΔ18 C-DI to AMPAR** Structural comparison of NaKΔ18 C-DI and GluA2. (a) Selectivity filter of NaKΔ18 C-DI with the $2F_o$-$F_c$ electron density contoured at 1σ. (b) Selectivity filter of GluA2 with the cryo-EM map contoured at 1σ (PDB ID 5weo). (c) Superposition of the selectivity filters of NaKΔ18 C-DI and GluA2. Only two diagonally opposed subunits are shown for clarity. (d) Comparison of the filter dimensions of NaKΔ19 (PDB ID 3E8H), NaKΔ18 C-DI and GluA2.

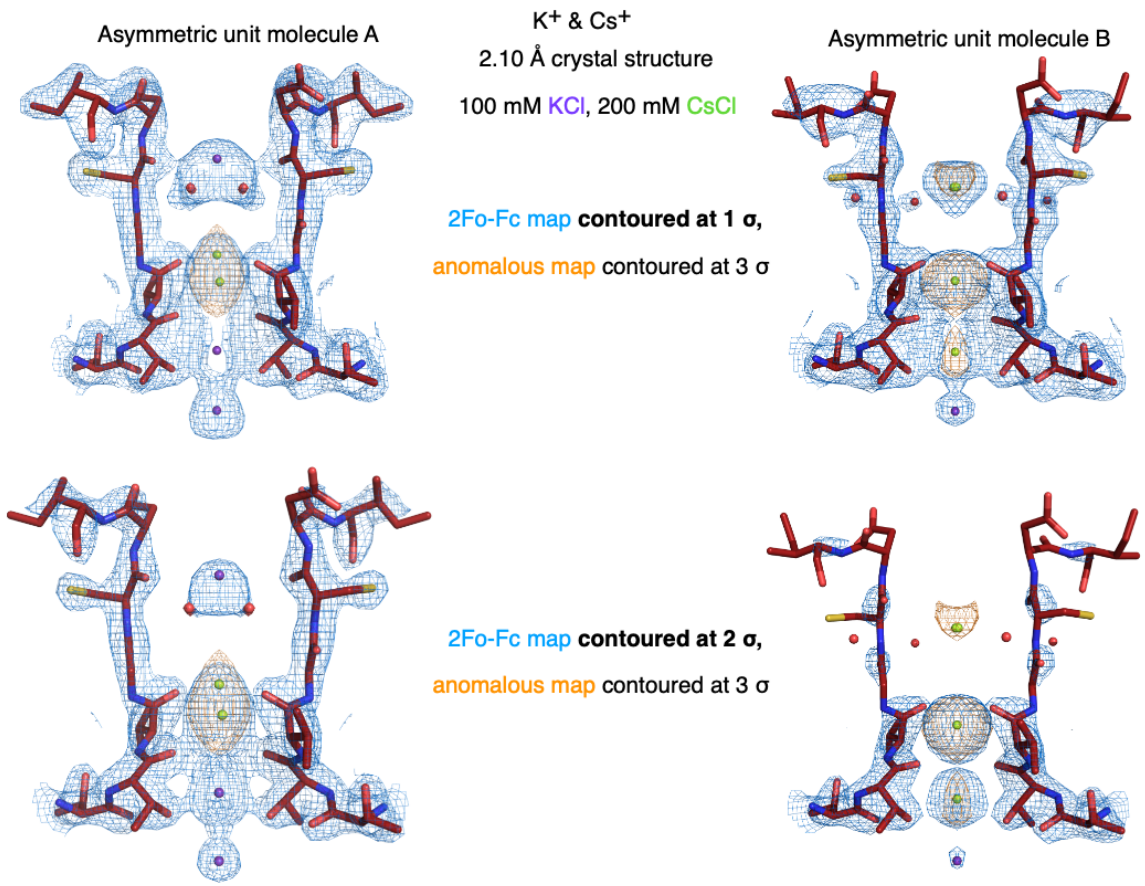


**Supplementary Figure 2 Ion occupancy varies by molecule in the asymmetric unit.** The $Cs^{+}$ complex crystallised in space group I4 and contains two subunits in the asymmetric unit from individual tetramers. (a) In tetramer A, $K^{+}$ occupies binding site $S_4$ and the vestibule along with water molecules. $Cs^{+}$ exists in two alternative conformations in binding site $S_3$. (b) In tetramer B, $Cs^{+}$ displaced $K^{+}$ from all binding sites within the filter. In both tetramers, one $K^{+}$ ion sits at the intracellular entrance to the selectivity filter. The $2F_o$-$F_c$ electron density (blue mesh) is contoured at 1σ (upper) and 2σ (lower) to highlight the less well refined structure of asymmetric unit molecule B. The anomalous signal of $Cs^{+}$ (orange mesh) is contoured at 3σ.

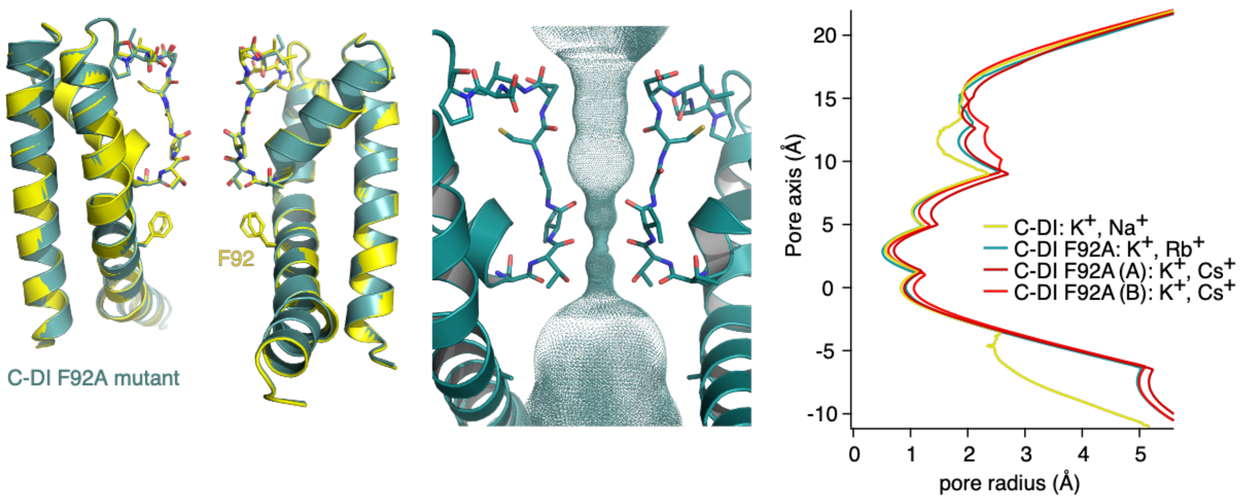


**Supplementary Figure 3 Effect of F92A mutation on the ion conduction pathway.** (left) Structural alignment of NaKΔ18 C-DI (yellow) with NaKΔ18 C-DI F92A (teal). (middle) Solvent-accessible pathway of NaKΔ18 C-DI F92A calculated with Hole, showing also the large cavity underneath the selectivity filter. (right) Pore profiles of NaKΔ18 C-DI ($Na^+$ and $K^+$) and NaKΔ18 C-DI F92A ($K^+$ and either $Rb^+$ or $Cs^+$).

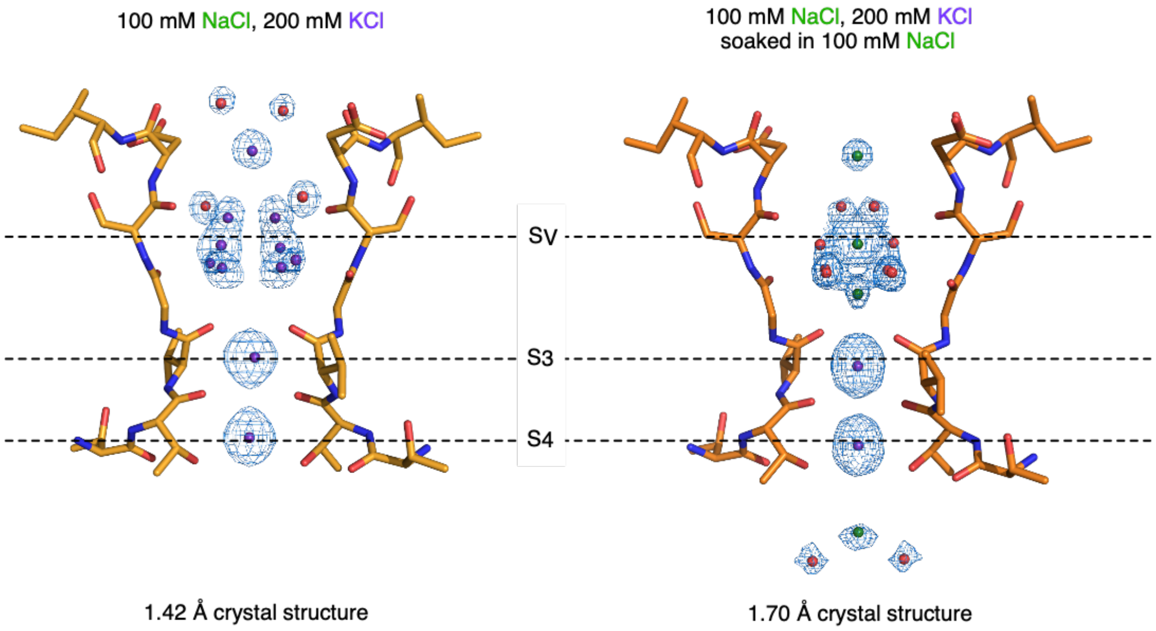


**Supplementary Figure 4 $Na^+$ binding in NaKΔ18 S-DI.** $K^+$ cannot be replaced by $Na^+$ in binding sites $S_3$ and $S_4$ even after prolonged soaking in NaCl solution. In contrast, $Na^+$ ions are evident in the vestibule site ($S_v$).

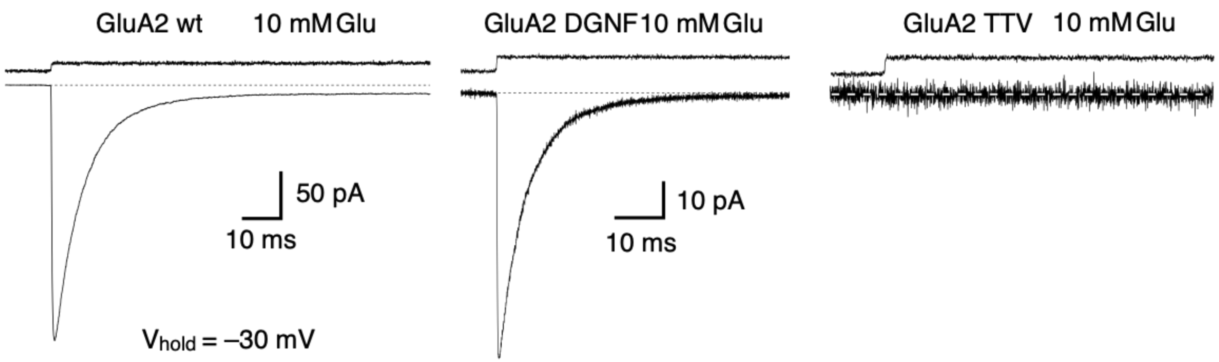


**Supplementary Figure 5 Patch-clamp recordings of GluA2 mutants.** (left) Macroscopic current in outside out patches from HEK-293 cells of WT GluA2 in response to 10 mM glutamate. (middle) Mutating C-DI stretch in GluA2 to DGNF has little effect on the current response to glutamate. (right) Including the pore loop mutation GluA2 TTV completely eliminates the response to 10 mM glutamate.

at two
2 and
and
values
error
field,

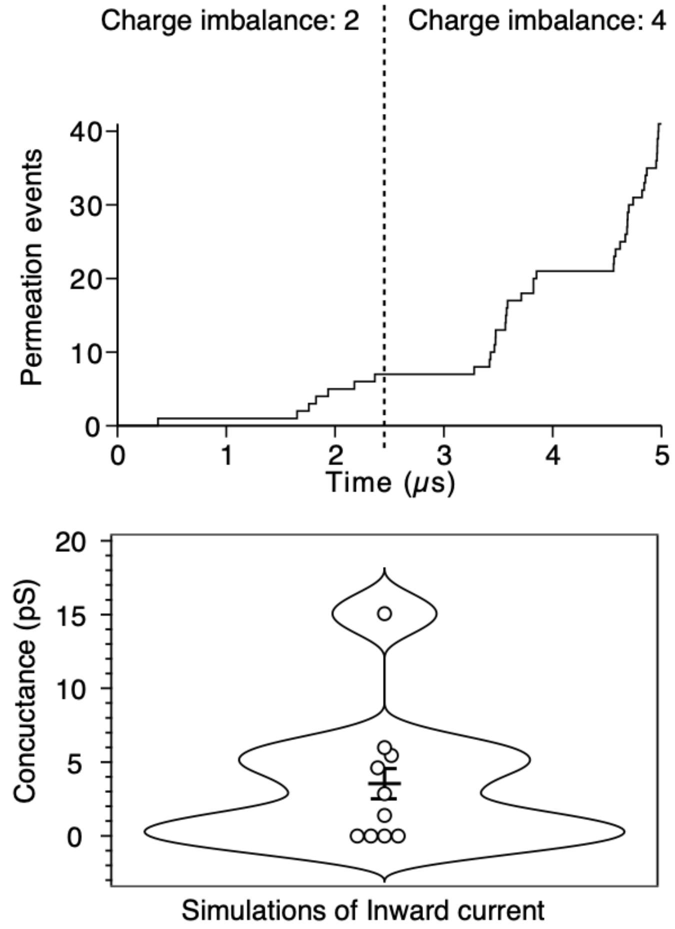


**Supplementary Figure 6 Potassium conductance in NaKΔ18 C-DI derived from CHARMM36m simulations.** (upper) Cumulative inward permeation events versus time different transmembrane voltages. Charge imbalances of 4 correspond to transmembrane voltages of 220±10 mV 420±10 mV, respectively. (lower) Simulated conductance for each of the individual 500 ns trajectories. Mean of the conductance is shown as a horizontal line with standard of the mean. In the simulations with Charmm36m force only inward $K^+$ ion permeation was observed.

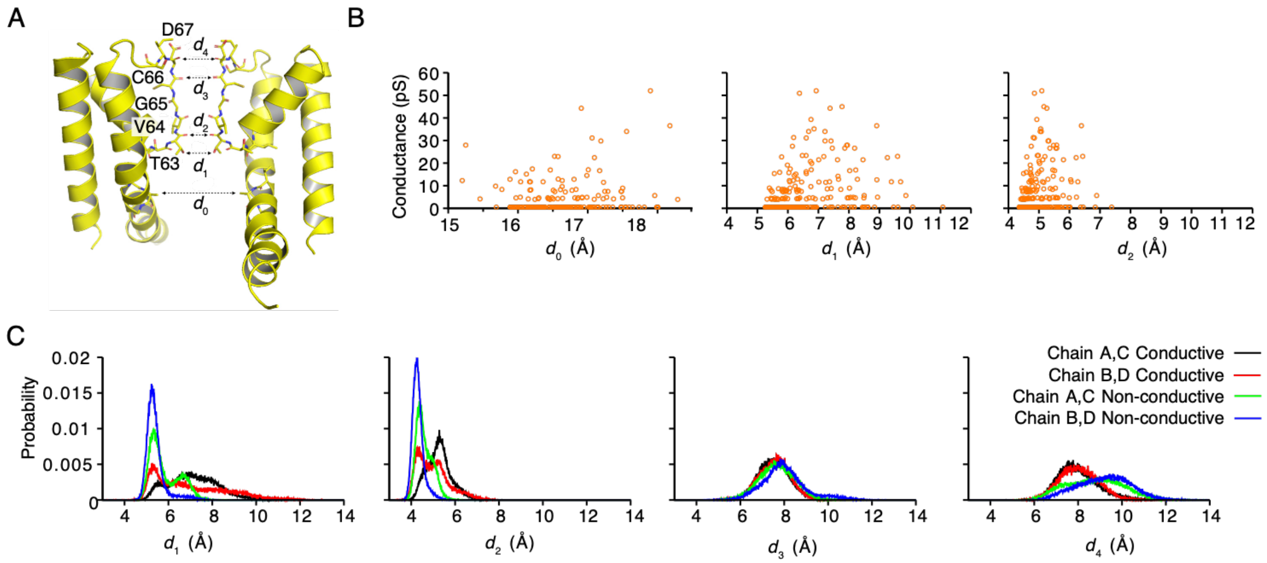


**Supplementary Figure 7 Diameter of the SF and $K^+$ ion conductance.** (A) illustration of distances $d_0$-$d_4$ in the NaK C-DI channel. (B) $K^+$ conductance (orange circles) as a function of distances $d_0$, $d_1$ and $d_2$. (C) Comparison between conductive and non-conductive trajectories for the distribution of distances $d_1$, $d_2$, $d_3$ and $d_4$ between residues in chain A-C and chain B-D sampled during the simulations. The simulations were performed with AMBER99SB force field.

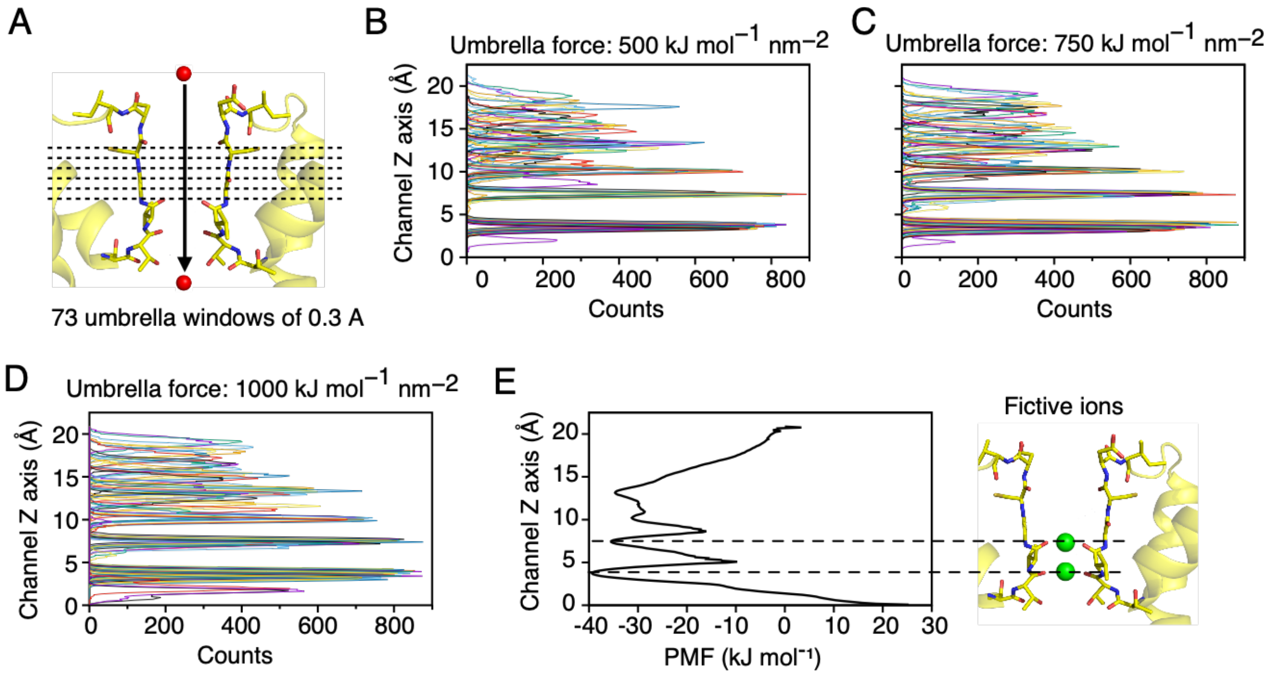


**Supplementary Figure 8 Umbrella sampling for sodium conduction in NaKΔ18 C-DI.** (A) A schematic representation of the pulling simulation, where a single $Na^+$ ion was pulled through the central axis of the selectivity filter in the NaK C-DI channel. (B-D) Histograms of overlaid umbrella sampling windows for simulations with three different harmonic biasing potentials (spring constants of 500, 750 and 1000 kJ $mol^{-1}$ $nm^{-2}$). (E) One-dimensional potential of mean force across the NaK C-DI pore aligned with (f) the X-ray structure of NaK C-DI showing two fictive $Na^+$ (green balls) at two main binding sites in the SF. The umbrella sampling simulations were performed with CHARMM36m force field.

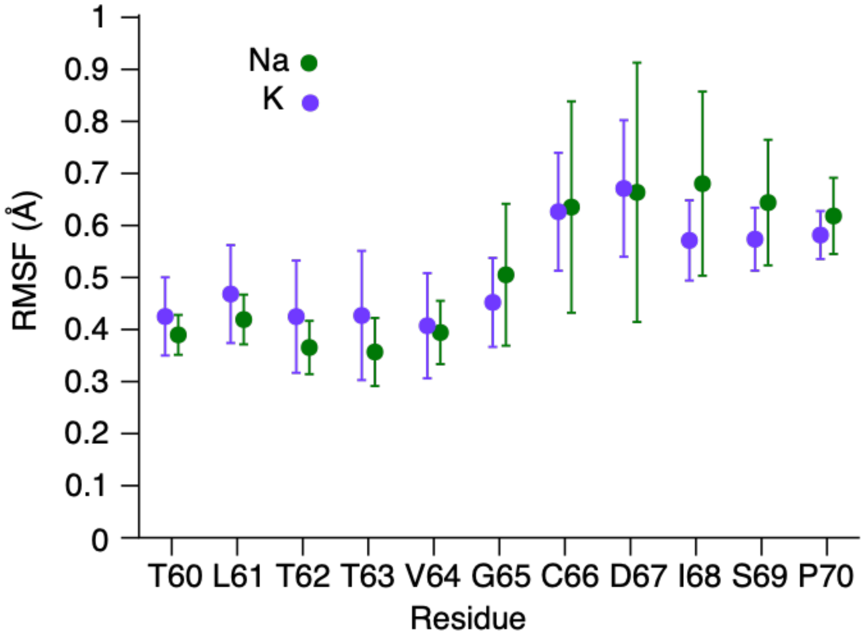


**Supplementary Figure 9 RMSF for the selectivity filter.** Comparison of the RMSF values in the SF region for simulations with $K^+$ (purple) and $Na^+$ (green). The simulations were performed with AMBER99sb force field.

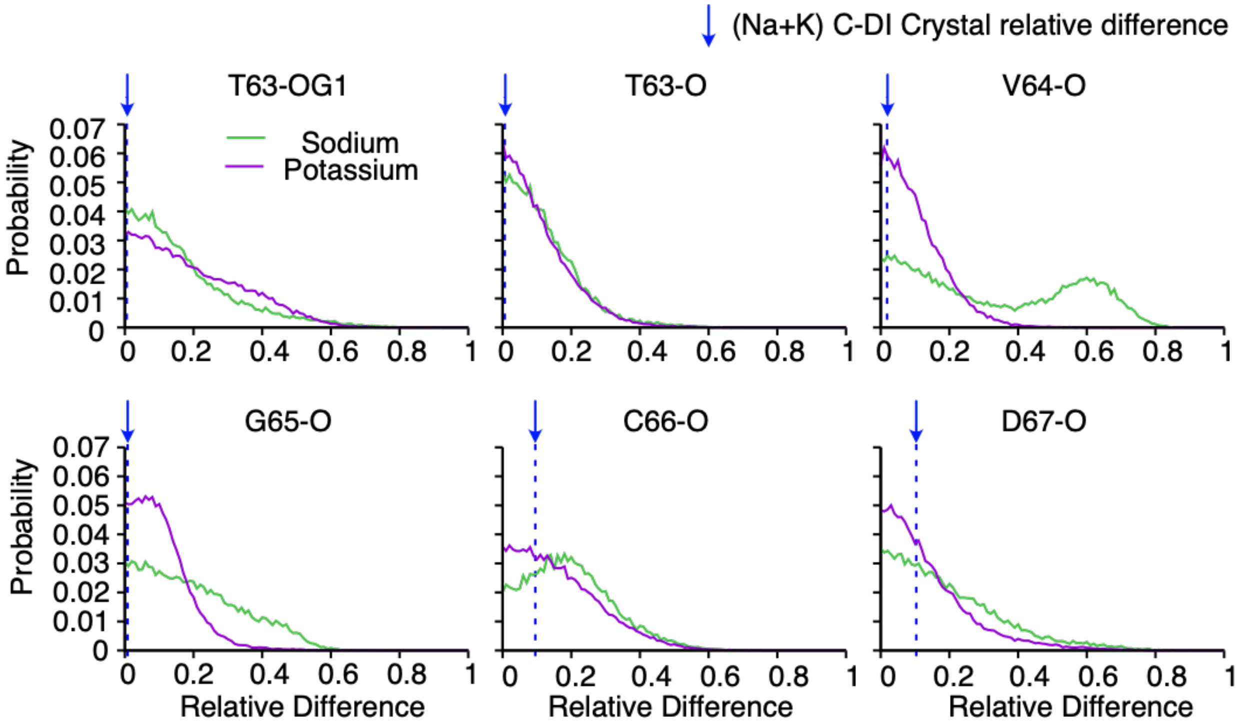


**Supplementary Figure 10 Analysis of SF asymmetry from MD simulations.** Comparison of the relative difference (AB,BC) distributions sampled from simulations with $K^+$ (purple) and $Na^+$ (green). The relative difference parameter was calculated for (b-f) the backbone oxygen in the SF of the NaK C-DI and (a) the side chain oxygen of T63. The vertical blue dashed lines represent the relative difference (AB,BC) values of the crystal structure.

**Supplementary Table 1.** Crystallography data collection and refinement statistics.

**Table 1. Data collection and refinement statistics.**

| | **C-DI $Na^+/K^+$ 7OOR** | **C-DI $Li^+/K^+$ 7OOU** | **C-DI F92A $Rb^+/K^+$ 7PA0** | **C-DI F92A $Cs^+/K^+$ 8A7X** | **S-DI $Na^+/K^+$ 7OPH** | **S-ELM $Na^+/K^+$ 7OQ1** | **S-DI Na+ soak 7OQ2** | **C-DI $Rb^+/Na^+$ 8A35** |
|---|---|---|---|---|---|---|---|---|
| **Wavelength** | 0.9184 | 0.9184 | 0.8154 | 0.918 | 1.033 | 1.033 | 1.033 | 0.815 |
| **Resolution range** | 40.62 - 1.47 (1.523 - 1.47) | 38.2 - 1.796 (1.86 - 1.796) | 40.44 - 1.95 (2.02 - 1.95) | 45.12 - 2.1 (2.175 - 2.1) | 38.01 - 1.418 (1.469 - 1.418) | 59.45 - 1.85 (1.916 - 1.85) | 59.73 - 1.7 (1.761 - 1.7) | 32.92 - 2.05 (2.123 - 2.05) |
| **Space group** | C 2 2 21 | C 2 2 21 | C 2 2 21 | I 4 | C 2 2 21 | C 2 2 21 | C 2 2 21 | I 4 |
| **Unit cell** | 81.25 88.49 49.66 90 90 90 | 81.481 88.413 49.586 90 90 90 | 80.874 87.312 49.105 90 90 90 | 68.16 68.16 90.25 90 90 90 | 81.13 88.021 49.313 90 90 90 | 81.214 87.277 48.98 90 90 90 | 81.204 88.173 49.389 90 90 90 | 68.062 68.062 90.268 90 90 90 |
| **Total reflections** | 522176 (41899) | 110894 (10285) | 63802 (5721) | 50694 (5160) | 214951 (20474) | 30031 (2596) | 39646 (3919) | 58263 (5884) |
| **Unique reflections** | 30700 (2990) | 17002 (1616) | 23961 (1231) | 22957 (2360) | 33738 (3284) | 15103 (1357) | 19895 (1966) | 24514 (1276) |
| **Multiplicity** | 17.0 (14.0) | 6.5 (6.4) | 2.7 (2.5) | 2.2 (2.2) | 6.4 (6.2) | 2.0 (1.9) | 2.0 (2.0) | 2.4 (2.4) |
| **Completeness (%)** | 99.52 (98.81) | 99.30 (95.73) | 98.86 (95.33) | 96.07 (96.97) | 99.60 (97.96) | 99.06 (91.07) | 99.80 (99.69) | 99.71 (99.77) |
| **Mean** | 17.82 (1.20) | 12.42 (0.95) | 5.56 (0.72) | 44.22 (1.42) | 12.64 (0.71) | 11.46 (1.60) | 4.46 (0.60) | 12.43 (1.01) |

| | | | | | | | | |
|---|---|---|---|---|---|---|---|---|
| **I/sigma(I)** | | | | | | | | |
| **Wilson B-factor** | 20.10 | 27.97 | 34.14 | 44.36 | 21.55 | 24.96 | 21.96 | |
| **R-merge** | 0.1012 (2.622) | 0.1172 (1.913) | 0.1291 (1.443) | 0.1228 (0.9152) | 0.08321 (2.57) | 0.03287 (0.4445) | 0.06674 (0.9783) | 0.04077 (0.8466) |
| **R-meas** | 0.1044 (2.722) | 0.1275 (2.082) | 0.1605 (1.821) | 0.1558 (1.168) | 0.09071 (2.805) | 0.04648 (0.6286) | 0.09439 (1.384) | 0.05157 (1.066) |
| **R-pim** | 0.02517 (0.7174) | 0.04932 (0.8092) | 0.09417 (1.095) | 0.09434 (0.7146) | 0.03569 (1.108) | 0.03287 (0.4445) | 0.06674 (0.9783) | 0.03111 (0.6391) |
| **CC1/2** | 0.999 (0.44) | 0.999 (0.351) | 0.995 (0.426) | 0.936 (0.392) | 0.999 (0.376) | 0.999 (0.693) | 0.996 (0.596) | 0.999 (0.41) |
| **CC*** | 1 (0.781) | 1 (0.721) | 0.999 (0.773) | 0.983 (0.75) | 1 (0.739) | 1 (0.905) | 0.999 (0.864) | 1 (0.763) |
| **Reflections used in refinement** | 30695 (2988) | 16993 (1614) | 12892 (1225) | 22848 (2336) | 33697 (3273) | 15100 (1357) | 19862 (1961) | 12887 (1276) |
| **Reflections used for R-free** | 1535 (149) | 849 (80) | 643 (61) | 1139 (113) | 1683 (163) | 1516 (136) | 1969 (178) | 645 (64) |
| **R-work** | 0.1477 (0.3404) | 0.1767 (0.3843) | 0.2050 (0.3258) | 0.2010 (0.3260) | 0.1535 (0.3378) | 0.1600 (0.2490) | 0.1860 (0.3211) | 0.1874 (0.3202) |
| **R-free** | 0.1850 (0.4223) | 0.2100 (0.4062) | 0.2456 (0.3467) | 0.2451 (0.4421) | 0.1808 (0.3807) | 0.1858 (0.2832) | 0.2333 (0.3485) | 0.2157 (0.3185) |
| **CC(work)** | 0.966 (0.762) | 0.962 (0.679) | 0.948 (0.596) | 0.896 (0.586) | 0.965 (0.690) | 0.955 (0.870) | 0.959 (0.691) | 0.850 (0.371) |
| **CC(free)** | 0.945 (0.650) | 0.933 (0.632) | 0.965 (0.802) | 0.920 (0.561) | 0.966 (0.629) | 0.976 (0.770) | 0.947 (0.602) | 0.817 (0.428) |
| **Number of** | 1755 | 1672 | 1648 | 1635 | 1742 | 1663 | 1718 | 1490 |

| | | | | | | | | |
|---|---|---|---|---|---|---|---|---|
| **non-hydrogen atoms** | | | | | | | | |
| **macromolecules** | 1532 | 1472 | 1455 | 1453 | 1514 | 1503 | 1469 | 1409 |
| **ligands** | 427 | 347 | 401 | 406 | 384 | 266 | 404 | 162 |
| **solvent** | 62 | 54 | 44 | 28 | 82 | 42 | 97 | 17 |
| **Protein residues** | 186 | 185 | 184 | 187 | 187 | 187 | 186 | 181 |
| **RMS(bonds)** | 0.007 | 0.008 | 0.003 | 0.014 | 0.008 | 0.013 | 0.011 | 0.007 |
| **RMS(angles)** | 0.86 | 0.96 | 0.54 | 1.43 | 0.98 | 0.97 | 0.96 | 0.54 |
| **Ramachandran favored (%)** | 98.90 | 100.00 | 98.89 | 98.91 | 99.45 | 99.45 | 99.45 | 96.61 |
| **Ramachandran allowed (%)** | 1.10 | 0.00 | 1.11 | 1.09 | 0.55 | 0.55 | 0.55 | 3.39 |
| **Ramachandran outliers (%)** | 0.00 | 0.00 | 0.00 | 0.00 | 0.00 | 0.00 | 0.00 | 0.00 |

| | | | | | | | | |
|---|---|---|---|---|---|---|---|---|
| **Rotamer outliers (%)** | 0.00 | 0.59 | 0.00 | 0.60 | 0.57 | 0.00 | 0.00 | 0.62 |
| **Clashscore** | 5.06 | 4.16 | 4.72 | 10.33 | 4.89 | 2.69 | 3.22 | 21.02 |
| **Average B-factor** | 28.11 | 31.43 | 41.62 | 63.29 | 29.98 | 32.50 | 31.23 | 57.65 |
| **macromolecules** | 24.14 | 28.68 | 40.07 | 60.77 | 26.83 | 30.52 | 28.12 | 56.62 |
| **ligands** | 61.16 | 54.73 | 54.52 | 86.87 | 54.48 | 54.74 | 55.70 | 79.07 |
| **solvent** | 40.33 | 43.28 | 49.17 | 64.21 | 44.43 | 40.67 | 39.92 | 61.88 |

Statistics for the highest-resolution shell are shown in parentheses.

**Supplementary Table 2** Single channel kinetics of NaK C-DI F92A mutant in bilayers

| Closed | mean | *N* | Open | mean | *N* |
|---|---|---|---|---|---|
| time constants | | | | | |
| $\tau_1$ (ms) | 2012 ± 595 | 3 | | 155 ± 42 | 3 |
| $\tau_2$ (ms) | 234 ± 71 | 3 | | 35 ± 8 | 4 |
| $\tau_3$ (ms) | 63 ± 28 | 4 | | 6 ± 0.4 | 3 |
| $\tau_4$ (ms) | 11 ± 6 | 4 | | | |
| amplitudes | | | | | |
| $a_1$ (%) | 21 ± 10 | 4 | | 16 ± 9 | 4 |
| $a_2$ (%) | 17 ± 6 | 4 | | 51 ± 16 | 4 |
| $a_3$ (%) | 30 ± 10 | 4 | | 33 ± 12 | 4 |
| $a_4$ (%) | 33 ± 11 | 4 | | | |

The number of values averaged is indicated. Four recordings were used, for several recordings there were fewer than 4 or 3 components in the shut or open time exponential distributions. For these, we included the amplitude of zero in the average.

**Supplementary Table 3.** Summary of MD-based computational electrophysiology simulations of the NaK C-DI channel for $K^+$ and $Na^+$ using two different force fields (Amber: AMBER99sb; Charmm: CHARMM36) and varying transmembrane potentials. All simulations were carried out with the TIP3P water model at a temperature of 310 K using an ion concentration of 600 mM. Errors are the standard deviations.

| Ion | Force field | Starting Structure | Charge imbalance | Potential (mV) | Number of 500 ns simulations | Total simulation time ($\mu$s) | Direction of ion flow | Ion permeation event | Conductance (pS) |
|---|---|---|---|---|---|---|---|---|---|
| $K^+$ | Amber | NaK C-DI with $Rb^+$ | 2 | 180 ± 50 | 36 | 18 | Inward | 104 | 4.5 ± 5.1 |
| | | | | | | | Outward | 45 | 2.9 ± 4.5 |
| | | | 4 | 410 ± 40 | 33 | 16.5 | Inward | 223 | 5.2 ± 3.9 |
| | | | | | | | Outward | 262 | 6.4 ± 6.2 |
| | Charmm | NaK C-DI with $Rb^+$ | 2 | 220 ± 10 | 10 | 5 | Inward | 7 | 1.9 ± 2.0 |
| | | | | | | | Outward | 0 | 0 |
| | | | 4 | 420 ± 10 | 10 | 5 | Inward | 34 | 5.1 ± 5.5 |
| | | | | | | | Outward | 1 | 0 |
| $Na^+$ | Amber | NaK C-DI with $Na^+$ + $K^+$ | 2 | 180 ± 20 | 25 | 12.5 | Inward | 0 | 0 |
| | | | | | | | Outward | 0 | 0 |
| | | | 4 | 430 ± 30 | 25 | 12.5 | Inward | 0 | 0 |
| | | | | | | | Outward | 6 | 0.2 ± 0.4 |
| | Charmm | NaK C-DI with $Na^+$ + $K^+$ | 2 | 200 ± 10 | 10 | 5 | Inward | 0 | 0 |
| | | | | | | | Outward | 0 | 0 |
| | | | 4 | 410 ± 10 | 10 | 5 | Inward | 0 | 0 |
| | | | | | | | Outward | 0 | 0 |
